# SOLAR Mediates Lysosomal Membrane Repair and Redox Defense

**DOI:** 10.64898/2026.07.08.736954

**Authors:** Jinrui Xun, Zhenjie Yi, Haoxiang Yang, Jenny Cheng, William A. Dion, Rae Yang, Xiaojie Yu, Fangling Tian, Bo Lv, Henry F. Liu, Anuj K. Peri, Bokai Zhu, Jay Xiaojun Tan

## Abstract

Lysosomal damage is increasingly recognized as a hallmark of aging. Understanding lysosomal quality control may open new therapeutic strategies to enhance lysosomal resilience. Here, we identify SOLAR (<u>S</u>QSTM1/p62 <u>O</u>ligomer-mediated <u>L</u>ysosomal <u>A</u>ntioxidant Defense And Membrane <u>R</u>epair), a lysosomal quality control pathway that links CASM (conjugation of ATG8 to single membranes) to p62-driven membrane repair and redox signaling. In this pathway, CASM, but not macroautophagy, recruited p62 to damaged lysosomes. Efficient lysosomal recruitment of p62 required its ATG8-binding and self-oligomerizing domains, enabling p62 assemblies to promote membrane repair. p62 mutations associated with neurodegeneration interfered with p62 recruitment and compromised lysosomal repair. Besides membrane repair, the SOLAR pathway also activated the p62-KEAP1-NRF2 redox signaling axis, driving transcriptional upregulation of antioxidative genes and cholesterol biosynthesis. By integrating membrane repair and redox defense, SOLAR establishes a coordinated lysosomal quality control pathway with implications for aging and degenerative disease.

## INTRODUCTION

Lysosomes are essential subcellular organelles with critical roles in cellular stress clearance and organismal health. As the general organelles degrading diverse cellular cargos including protein aggregates and invading pathogens, lysosomes are prone to membrane damage, known as lysosomal membrane permeabilization (LMP)^1–3^. In aging and diseases, persistent lysosomal damage and repair failure are emerging as key mechanisms driving pathogenesis^1^. A comprehensive understanding of lysosomal quality control mechanisms could pave the way for novel strategies in disease prevention and therapy.

In healthy cells, multiple quality control pathways are activated by LMP to repair, remove, and replace damaged lysosomes^1–4^. The transcription factor EB (TFEB) and TFE3 pathway is a major transcriptional driver of stress-induced lysosomal biogenesis^5,6^, whereas lysophagy mediates the turnover of overly damaged lysosomes^7–9^. Multiple additional pathways are activated by LMP to mediate rapid lysosomal membrane repair, such as membrane involution by the endosomal sorting complex required for transport (ESCRT) pathway^10,11^ or sphingomyelin-to-ceramide transition^12^ and membrane patching using lipids through the phosphoinositide-initiated membrane tethering and lipid transport (PITT) pathway^13,14^, as well as membrane stabilization by protein assemblies such as annexins^15,16^ and stress granule^17,18^. There are also potential roles of membrane tubulation in the removal of damaged lysosomal membranes and lysosomal reformation^19–21^. It remains unclear to what extent each pathway contributes to lysosomal repair and how such diverse repair mechanisms are coordinated.

One hallmark response of lysosomal damage is the direct conjugation of ATG8s (LC3s and GABARAPs) onto lysosomal membranes^22^. Such Atg8ylation is also known as conjugation of ATG8 to single membranes (CASM) or noncanonical autophagy, which happens independently of macroautophagy^22–24^. LMP-induced CASM appeared dramatic, as LC3 and GABARAP were almost completely lipidated in response to continuous lysosomal damage (**Figure S1A, B, C**). The level of Atg8ylation in response to LMP was much stronger than that observed during macroautophagy induction by nutrient starvation (Hank’s Balanced Salt Solution starvation or HBSS 1 h) or rapamycin and Torin1, inhibitors of mechanistic Target Of Rapamycin (mTOR) (**Figure S1A, B, C**). These observations suggest that LMP-induced CASM may serve as an essential platform for lysosomal quality control or cellular adaptations to lysosomal damage. Indeed, one established example is that CASM provides a key mechanism for TFEB/TFE3 activation to upregulate lysosomal biogenesis^25–29^. We therefore further explored how LMP-induced CASM mediates lysosomal damage response.

## RESULTS

### Identification of p62 as a potential effector of lysosomal Atg8ylation

We reasoned that the functions of LMP-induced CASM should be defined by effector proteins recruited to damaged lysosomes by ATG8s. To search for lysosomal CASM effectors, we compared the surface proteome of damaged lysosomes^14^ with established ATG8-binding proteins from BioGRID^30^. Among the resulting hits, p62 (also known as Sequestosome1 or SQSTM1) stood out as a top candidate based on the level of evidence supporting candidate-ATG8 interactions, and it was the only protein annotated to interact with all ATG8-family members (**Figure 1A, S1D, Supplementary Table S1**). We confirmed by immunofluorescence that endogenous p62 was robustly recruited to lysosomes in response to LMP across multiple cell types originated from distinct tissues, such as kidney (293T), cervical (HeLa), lung (Beas-2B or B2B) and bone (U2OS) (**Figure S1E-H**), which was further validated using multiple lysosomal markers (**Figure S1H-J**), as well as in iPSC-induced neurons (**Figure S1K**).

**Figure 1.**
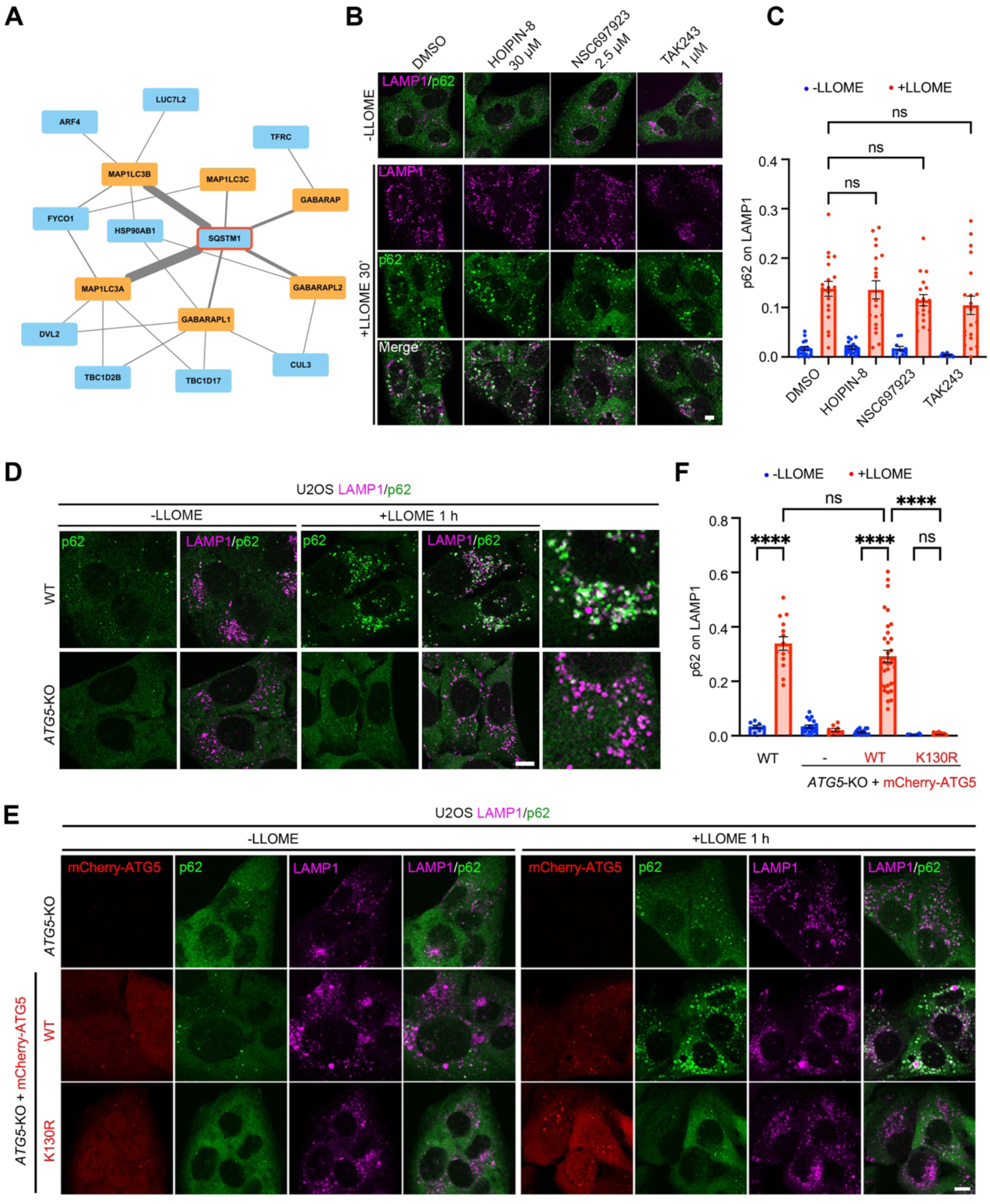
Identification of p62 as a potential effector of ATG8ylation but not ubiquitination on damaged lysosomes. (A) ATG8-interacting proteins enriched on damaged lysosomes. Protein-protein interactions were based on BioGRID database. Line thickness positively correlates to the levels of evidence. (B) p62 recruitment to damaged lysosomes is not affected by ubiquitination inhibition, as visualized by confocal immunofluorescence microscopy. U2OS cells were treated with DMSO or indicated concentrations of ubiquitination inhibitors for 2 h, and then treated with 500 μM of LLOME for 30 min, followed by fixation and immunostaining of endogenous LAMP1 and p62. (C) Quantification of p62 recruitment to lysosomes in (B). Data are means ± SEM; n = 17, 19, 19, 19, 11, 18, 10, 20 random cells, respectively. (D) p62 recruitment to damaged lysosomes is blocked in *ATG5*-KO cells, as visualized by immunofluorescence microscopy. WT or *ATG5*-KO U2OS cells were treated with vehicle or 500 μM of LLOME for 1 h, followed by fixation and immunostaining of endogenous LAMP1 and p62. (E) p62 recruitment to damaged lysosomes relies on ATG5-ATG12 conjugation. *ATG5* knockout cells or *ATG5* knockout cells stably expressing ATG5 WT or its K130R mutant with a mCherry-tag were treated with vehicle or 500 μM of LLOME for 1 h and then fixed for immunostaining of endogenous LAMP1 and p62. (F) Quantification of p62 recruitment to lysosomes in (F). Data are means ± SEM; n = 8, 14, 18, 8, 15, 34, 21, 44 random cells, respectively. Data were analyzed by two-way ANOVA with Tukey’s multiple comparison tests for (C) and (F). WT, wild type. KO, knockout. Bar, 10 μm. (G) See also Figure S1 and S2.

As an autophagy adaptor protein, p62 is most well known as an aggrephagy receptor^31–33^. In addition, p62 also mediates lysophagy, during which it is recruited to overly damaged lysosomes by ubiquitination^7,34^. However, pharmacologically eliminating cellular ubiquitination had a relatively modest impact on lysosomal recruitment of p62 following cell exposure to L-Leucyl-L-leucine methyl ester (LLOMe) (**Figure 1B, C, Figure S1L**), which mechanically damage lysosomes through fibril formation^35–37^. Given the strong lysosomal Atg8ylation following LMP (**Figure S1A, B, C**), we examined whether p62, as an ATG8-binding protein, was recruited by Atg8ylation. Remarkably, knockout of *ATG5*, an essential gene of the ubiquitination-like ATG8-conjugation system^38,39^, abolished LMP-induced lysosomal recruitment of p62 (**Figure 1D**). ATG5 contributes to ATG8ylation by functioning as the core catalytic scaffold of the E3-like ligase complex in which Lys130 (K130) of ATG5 is covalently conjugated to ATG12 (ref^40^). ATG5 K130R is a mutant losing ATG12 conjugation and is deficient in mediating ATG8 lipidation^40^. When reconstituted in *ATG5*-KO cells, wild-type ATG5, but not the K130R mutant, fully rescued p62 recruitment to damaged lysosomes (**Figure 1E, F, Supplementary Table S2**). *ATG5* knockout did not seem to affect LMP-induced ubiquitin puncta formation (**Figure S2A**). Therefore, the ATG8 conjugation activity, but not lysosomal ubiquitination, is essential for LMP-induced lysosomal recruitment of p62.

While p62 recruitment was completely gone in *ATG5* knockout cells within one hour of lysosomal damage by LLOMe, some weak p62 recruitment was detected after 3 h of continuous LLOMe treatment (**Figure S2B**). Such ATG5-independent p62 recruitment appeared to be ubiquitination-dependent, as the E1 inhibitor TAK243 which removes total cellular ubiquitination (**Figure S1L**) fully eliminated p62 recruitment in *ATG5*-KO cells (**Figure S2B**). Thus, Atg8ylation is a major mechanism driving p62 recruitment to damaged lysosomes, whereas ubiquitination mediates p62 recruitment at a later stage, probably to overly damaged lysosomes.

In contrast to p62 which was mainly recruited by Atg8ylation, the recruitment of other autophagy adaptors, such as OPTN, NDP52, TAX1BP1, and NBR1, were largely abrogated by the ubiquitination inhibitor TAK243 (**Figure S2C, D, E, F**), suggesting that ubiquitination on damaged lysosomes preferentially recruits autophagy adaptors other than p62. Thus, p62 is a unique adaptor protein that is recruited in an ATG5-dependent manner, suggesting distinct functions of p62 in lysosomal damage response compared with other adaptors.

### CASM, but not macroautophagy, recruits p62 to damaged lysosomes

Because ATG5 is essential for both CASM and macroautophagy, we carefully examined which function of ATG5 mediated p62 recruitment to damaged lysosomes. An E1 enzyme of the ATG8-conjugation system, ATG7, contributes to the conjugation of ATG12 to ATG5 (ref^39,41,42^). Similar to *ATG5*-KO, knockout of *ATG7* abolished p62 recruitment to damaged lysosomes (**Figure 2A**). In contrast, knockout of macroautophagy-specific genes not involved in CASM, such as *ATG2A/ATG2B*-DKO^14^, *ATG9A*-KO, and *ATG13*-KO, did not affect LMP-induced p62 recruitment (**Figure S3A-G**). These data strongly support a role for CASM but not macroautophagy in recruiting p62 to damage lysosomes.

**Figure 2.**
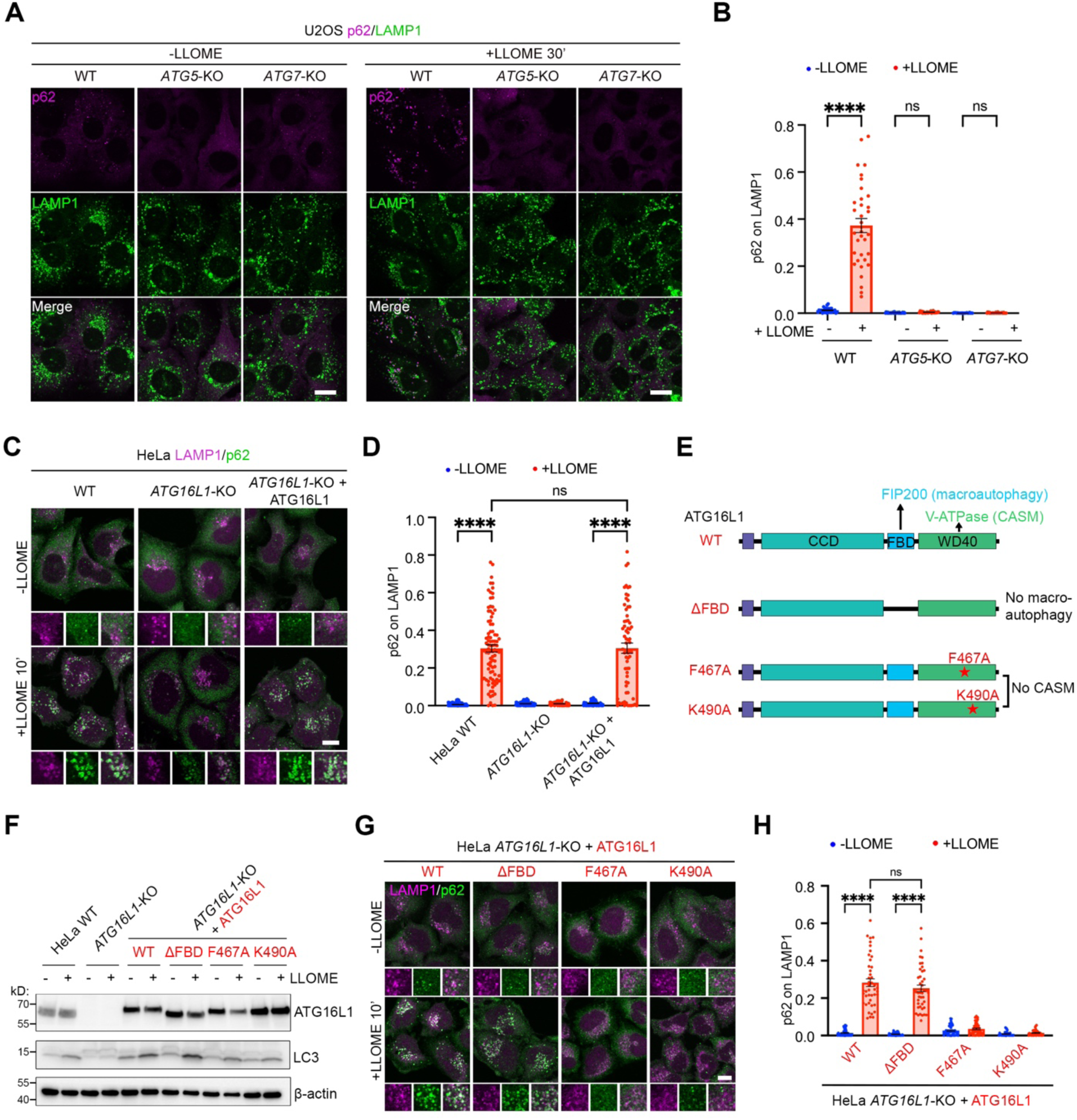
p62 is recruited to damaged lysosomes through CASM rather than macroautophagy. (A) *ATG5* or *ATG7* knockout abolishes p62 recruitment to damaged lysosomes, as visualized by confocal immunofluorescence microscopy. U2OS WT or *ATG5/7*-KO cells were treated with vehicle or 500 μM of LLOME for 30 min, followed by fixation and immunostaining of endogenous LAMP1 and p62. (B) Quantification of p62 recruitment to lysosomes in (A). Data are means ± SEM; n = 29, 35, 29, 26, 22, 33 random cells, respectively. (C) p62 recruitment to damaged lysosomes is abolished in *ATG16L1*-KO cells, which is rescued by reconstitution of *ATG16L1 in the knockout cells.* Indicated HeLa cells were treated with vehicle or 500 μM of LLOME for 10 min, followed by fixation and immunostaining of endogenous LAMP1 and p62. (D) Quantification of p62 recruitment to lysosomes in (C). Data are means ± SEM; n = 89, 92, 88, 83, 56, 68 random cells, respectively. (E) Illustration of ATG16L1 mutants generated to investigate the mechanism of p62 recruitment to damaged lysosomes. (F) Immunoblot showing the knockout and re-expression of *ATG16L1* and its mutants in HeLa cells. (G) p62 recruitment to damaged lysosomes requires the WD40 domain rather than FBD domain. HeLa *ATG16L1*-KO cells re-expressing ATG16L1 or its mutants were treated with vehicle or 500 μM of LLOME for 10 min, followed by fixation and immunostaining of endogenous LAMP1 and p62. (H) Quantification of p62 recruitment to lysosomes in (G). Data are means ± SEM; n = 45, 44, 38, 44, 44, 55, 32, 37 random cells, respectively. Data were analyzed by two-way ANOVA with Tukey’s multiple comparison tests for (B), (D) and (H). WT, wild type. KO, knockout. Bar, 10 μm. See also Figure S3.

During lysosomal damage, the ATG5–ATG12 complex can be recruited to lysosomes through either the V-ATPase-ATG16L1 axis or the TECPR1-ATG5 axis. The lysosomal proton pump V-ATPase recruits ATG16L1 upon the loss of lysosomal acidic pH^22,43^, whereas TECRP1 is recruited to lysosomes by LMP-induced sphingomyelin exposure on the outer leaflet of the lysosomal membrane^44–47^. Knockout of *ATG16L1* strongly reduced p62 recruitment, which was fully rescued by the reconstitution of wild-type ATG16L1 (**Figure 2C, D**), indicating that the V-ATPase-ATG16L1 axis is a major underlying mechanism.

ATG16L1 mediates CASM and macroautophagy by binding to V-ATPase and FIP200, respectively, through distinct domains^27^ (**Figure 2E**). We therefore further investigated whether the role of ATG16L1 in p62 recruitment was attributed to its activity in CASM or macroautophagy. In *ATG16L1*-KO cells, the reconstitution of an ATG16L1 mutant (ΔFBD), which loses FIP200 binding and therefore defective in macroautophagy, fully restored p62 recruitment to damaged lysosomes, similarly to the reconstitution of wild-type ATG16L1 (**Figure 2F-H** and **S3H**). In contrast, two ATG16L1 mutants (F467A or K490A) losing binding to V-ATPase were unable to rescue p62 recruitment (**Figure 2F-H** and **S3H**). Thus, the role of ATG16L1 in lysosomal CASM rather than macroautophagy contributes to LMP-induced p62 recruitment.

### LC3s and GABARAPs redundantly recruit p62

We next asked whether ATG8s themselves are necessary for p62 recruitment. In mammalian cells, there are six paralogs of ATG8: LC3A, LC3B, LC3C, GABARAP, GABARAPL1, and GABARAPL2, which have both redundant and distinct roles. While macroautophagy induction by either Torin1 or rapamycin triggered more puncta of LC3B than GABARAPL2, lysosomal damage by LLOME stimulated strong puncta of both (**Figure S1A** and **S1B**). Triple knockout of the *LC3*s (*LC3*-TKO) or *GABARAP*s (*GABARAP*-TKO)^48^ partially reduced p62 recruitment to damaged lysosomes (**Figure 3A, B, S4A**). Knockout of all six *ATG8*s (*ATG8*-HKO)^48^ largely blocked p62 recruitment (**Figure 3A, B, S4A**), indicating redundant roles of ATG8s in this pathway. Consistently, overexpression of RavZ, a *Legionella* effector that directly cleaves lipidated ATG8 from membranes, also strongly blocked lysosomal p62 recruitment (**Figure 3C, D**). In contrast, the enzymatic dead mutant of RavZ (C258A)^49^ did not affect p62 recruitment (**Figure 3C, D**). Thus, ATG8 proteins conjugated to the lysosomal membrane are essential for LMP-induced p62 recruitment.

**Figure 3.**
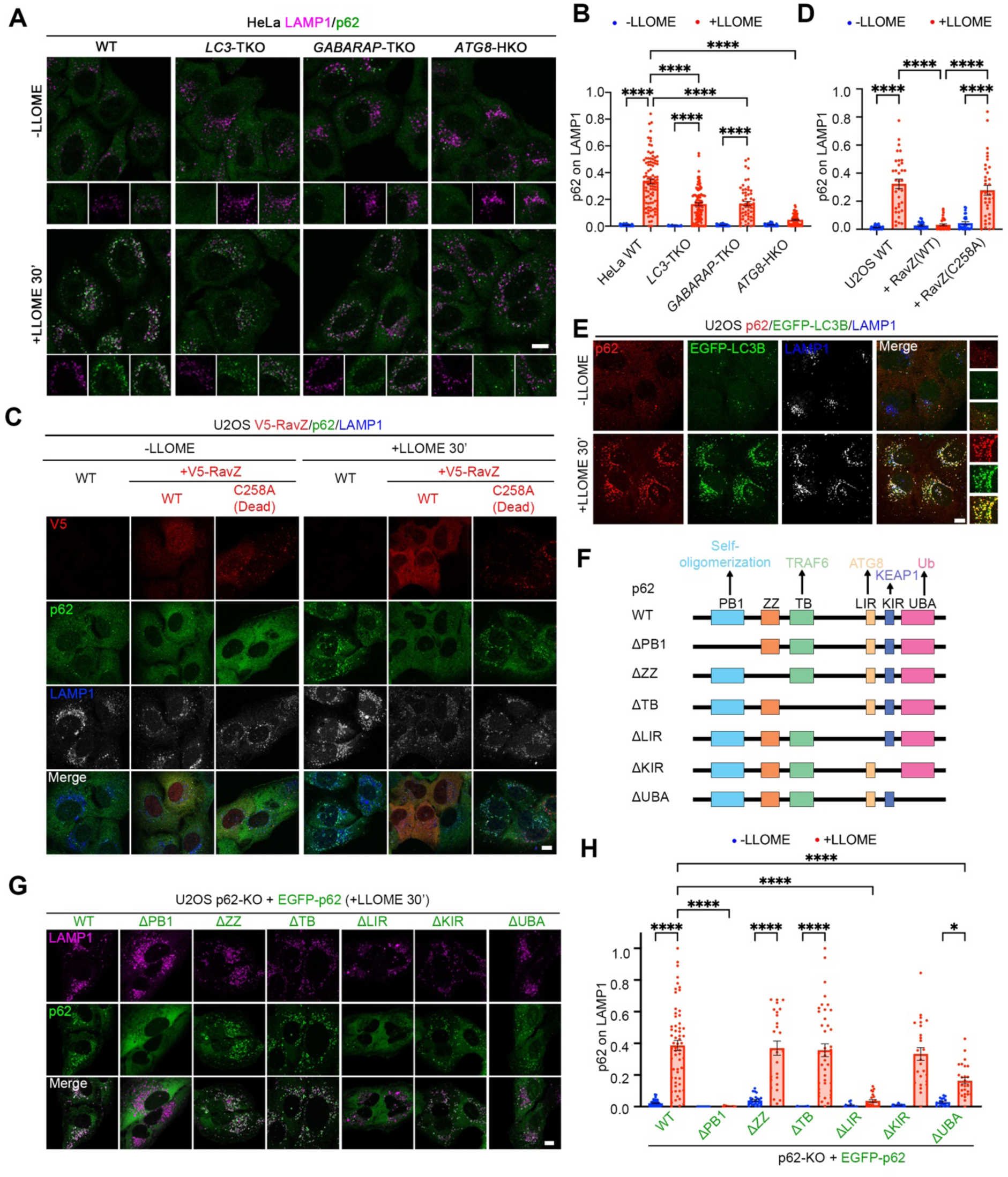
ATG8-binding and self-oligomerization mediate p62 recruitment to damaged lysosomes. (A) ATG8 family members, LC3s and GABARAPs, redundantly recruit p62 to damaged lysosomes. WT or indicated *ATG8*-knockout HeLa cells were treated with vehicle or 500 μM of LLOME for 30 min, followed by fixation and immunostaining of endogenous LAMP1 and p62. (B) Quantification of p62 recruitment to lysosomes in (A). Data are means ± SEM; n = 61, 97, 56, 96, 48, 57, 71, 73 random cells, respectively. (C) RavZ blocks p62 recruitment to damaged lysosomes. U2OS WT cells, with or without stable expression of V5-tagged RavZ or its catalytically dead mutant C258A, were treated with vehicle or 500 μM of LLOME for 30 min, followed by fixation and immunostaining of V5 tag and endogenous LAMP1 and p62. (D) Quantification of p62 recruitment to lysosomes in (C). Data are means ± SEM; n = 27, 34, 22, 36, 28, 36 random cells, respectively, pooled from three independent experiments. (E) LMP-induced p62 puncta strongly colocalize with LC3B. U2OS cells stably expressing EGFP-LC3B were treated with vehicle or 500 μM of LLOME for 30 min, followed by fixation and immunostaining of endogenous LAMP1 and p62. (F) Illustration of p62 domains and mutants generated to investigate the mechanism of its lysosomal recruitment. (G) p62 recruitment to damaged lysosomes requires its ATG8-binding LIR domain and self-oligomerization PB1 domain. U2OS p62-KO cells re-expressing EGFP-p62 or its truncation mutants were treated with 500 μM of LLOME for 30 min, followed by fixation and immunostaining of endogenous LAMP1 and p62. (H) Quantification of EGFP-p62 recruitment to lysosomes in (G). Data are means ± SEM; n = 55, 60, 18, 13, 21, 41, 14, 22, 41, 25, 17, 26 random cells, respectively, pooled from three independent experiments. Data were analyzed by two-way ANOVA with Tukey’s multiple comparison tests for (B), (D) and (H). WT, wild type. KO, knockout. Bar, 10 μm. See also Figure S4.

In line with a role for lysosomal Atg8ylation in p62 recruitment, LMP-induced p62 puncta largely colocalized with LC3 (**Figure 3E**). In contrast, LMP-induced ubiquitin puncta were restricted to a fraction of lysosomes that were mostly negative for LC3 (**Figure S4B, C**), and the ubiquitin puncta showed up at a later time point following lysosomal damage (**Figure S4D**). Furthermore, the lysosomal recruitment of p62 following the treatment with proton ionophores nigericin or monensin also relied on ATG5 (**Figure S4E**). Thus, lysosomal CASM appears to be a critical mechanism for p62 recruitment to stressed lysosomes.

### ATG8-binding and self-oligomerization drive stable lysosomal recruitment of p62

To further delineate the molecular basis of lysosomal p62 recruitment, we assessed the contribution of individual p62 domains to its localization. Based on AlphaFold-predicted structure^50^, four domains of p62 have defined structures: an N-terminal Phox and Bem1p (PB1) domain, a ZZ-type Zinc Finger (ZZ) domain, a TRAF6 Binding (TB) domain, and a C-terminal Ubiquitin-Associated (UBA) domain (**Figure 3F**). The four domains are connected by three flexible linkers, the longest of which is an intrinsically disordered region (IDR) located between the TB domain and the UBA domain (**Figure 3F**). An LC3-interacting region (LIR) was previous identified in this IDR^33^. We individually deleted each of the four structurally defined domains or the LIR motif to examine their impacts on p62 localization (**Figure 3F**).

When stably expressed in p62-KO cells, three of the deletion mutants, p62ΔZZ, ΔTB, and ΔUBA, were still substantially recruited to damaged lysosomes (**Figure 3G, H, and S4F**). Deletion of the LIR motif, which blocks p62 binding to ATG8 proteins, strongly suppressed p62 recruitment (**Figure 3G, H**). Notably, deletion of the PB1 self-oligomerization domain also completely eliminated p62 recruitment (**Figure 3G, H**). Therefore, ATG8 binding alone was insufficient for robust lysosomal recruitment of p62; instead, PB1-mediated self-oligomerization promotes recruitment by allowing multiple p62 molecules to engage membrane-conjugated ATG8 simultaneously.

### CASM mediates the recruitment of both p62 and ESCRT

We explored how cells coordinate the lysosomal recruitment of p62 along with components of established lysosomal membrane repair pathways, such as ESCRT^10,11^ and PITT^14,51^. Knockout of p62 did not seem to affect the lysosomal recruitment of PI4K2A, ORP9, or EGFP-ATG2A (**Figure S5A, B, C**), all of which are PITT pathway components^14^. Conversely, blocking the PITT pathway by *PI4K2A*-KO did not affect lysosomal recruitment of p62 (**Figure S5D**).

Knockout of p62 did not appear to affect LMP-induced lysosomal recruitment of the ESCRT-III subunit IST1 (**Figure 4A, B**). Remarkably, co-depletion of two ESCRT subunits TSG101 and ALIX, which blocks LMP-induced ESCRT-III assembly^10^, dramatically increased p62 recruitment to damaged lysosomes (**Figure 4C, D**). These results suggest that ESCRT subunits likely compete with p62 for ATG8 interactions on damaged lysosomes. Indeed, knockout of *ATG5* largely abolished the assembly of both ALIX and IST1, two distinct ESCRT subunits (**Figure 4E, F, G, H, S5E, F**). Reconstitution of wild-type ATG5, but not its conjugation-deficient mutant K130R, fully restored LMP-induced lysosomal recruitment of ALIX and IST1 (**Figure 4E, F, G, H, S5E, F**), consistent with the recently reported roles of ATG8 in facilitating ESCRT assembly on damaged lysosomes^52–54^. Thus, LMP-induced lysosomal CASM supports the assemblies of both ESCRT and p62 oligomers on lysosomal membranes.

**Figure 4.**
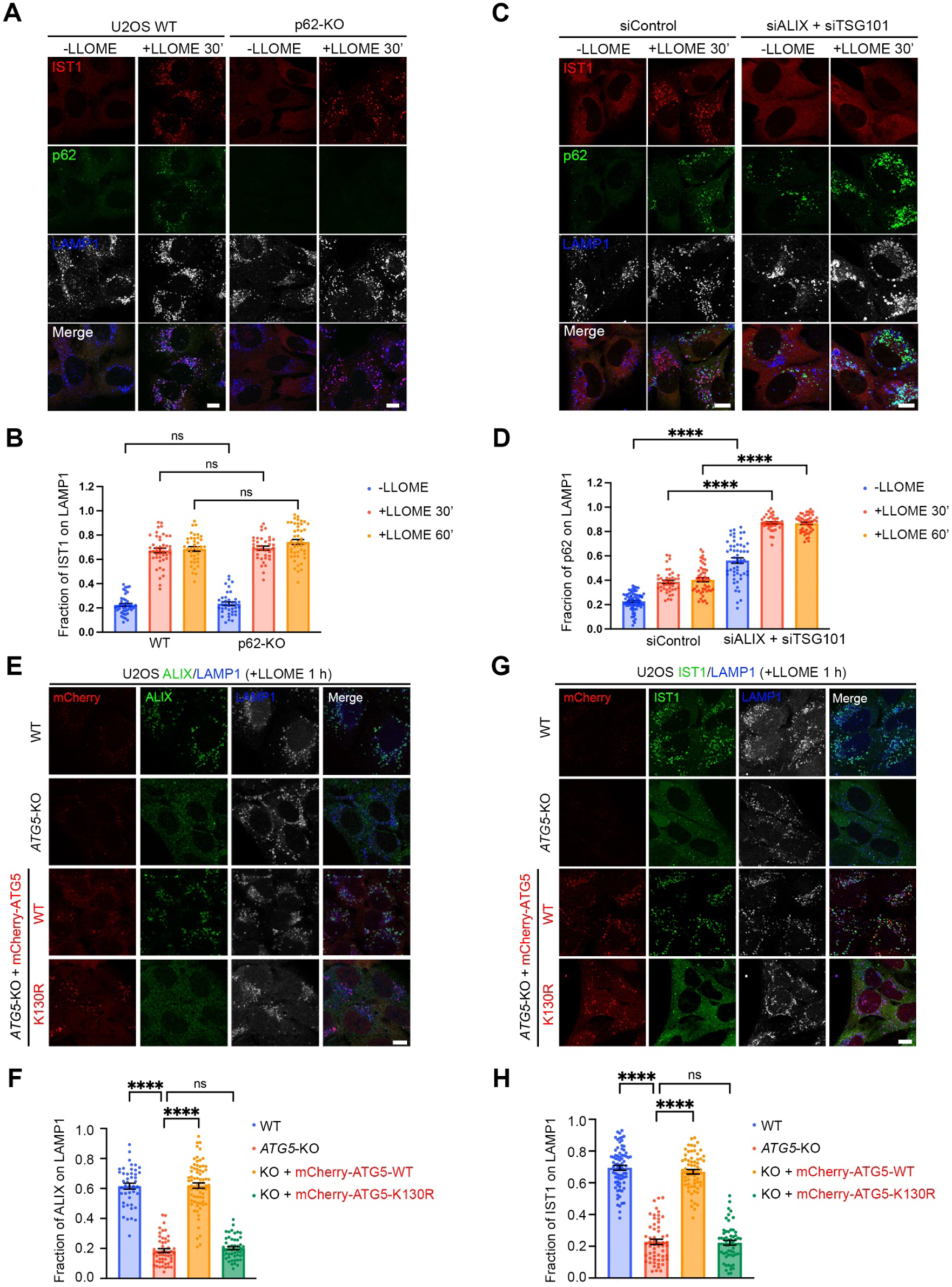
CASM mediates the recruitment of both p62 and ESCRT. (A) IST1 recruitment to damaged lysosomes is not affected by p62 knockout. U2OS WT or p62-KO cells were treated with vehicle or 500 μM of LLOME for 30 min, followed by fixation and immunostaining of endogenous IST1, LAMP1 and p62. (B) Quantification of the fraction of IST1 colocalized with LAMP1 in (A). Data are means ± SEM; n = 43, 44, 39, 38, 36, 46 random cells, respectively. (C) p62 recruitment to damaged lysosomes is dramatically increased by ESCRT knockdown, as visualized by immunofluorescence microscopy. U2OS cells were transfected with siRNAs against *ALIX*, *TSG101*, or a control siRNA, then treated with vehicle or 500 μM of LLOME for 30/60 min, followed by fixation and immunofluorescence staining of endogenous IST1, LAMP1 and p62. (D) Quantification of fraction of p62 colocalized with LAMP1/total p62 in (C). Data are means ± SEM; n = 76, 41, 49, 54, 38, 48 random cells, respectively. (E) ATG5 mediates ALIX recruitment to damaged lysosomes through its conjugation activity. U2OS WT, *ATG5*-KO cells or *ATG5*-KO cells re-expressing ATG5 or its conjugation-dead mutant K130R were treated with vehicle or 500 μM of LLOME for 1 h, followed by fixation and immunostaining of endogenous ALIX and LAMP1. (F) Quantification of the fraction of ALIX colocalized with LAMP1 in (E). Data are means ± SEM; n = 44, 50, 73, 50 random cells, respectively. (G) ATG5 mediates IST1 recruitment to damaged lysosomes through its conjugation activity. U2OS WT, *ATG5*-KO cells or *ATG5*-KO cells re-expressing ATG5 or its conjugation-dead mutant K130R were treated with vehicle or 500 μM of LLOME for 1 h, followed by fixation and immunostaining of endogenous IST1 and LAMP1. (H) Quantification of the fraction of IST1 colocalized with LAMP1 in (G). Data are means ± SEM; n = 76, 57, 65, 56 random cells, respectively. Data were analyzed by two-way (B and D) and one-way (F and H) ANOVA with Tukey’s multiple comparison tests. WT, wild type. KO, knockout. Bar, 10 μm. See also Figure S5.

### p62 assemblies mediate lysosomal membrane repair

We noted that both endogenous and reconstituted p62 proteins were stabilized following lysosomal damage (**Figure 5A**), suggesting that p62 is post-translationally stabilized on the lysosomal membrane. Consistently, all the p62 deletion mutants retaining their lysosomal recruitment capacity, including ΔKIR, ΔUBA, ΔZZ, and ΔTB (**Figure 3G and H**), were stabilized following lysosomal damage by LLOME (**Figure 5A**). By contrast, two mutants of p62, ΔPB1 and ΔLIR, which cannot be recruited to lysosomes (**Figure 3G and H**), were not stabilized (**Figure 5A**). Thus, the ATG8-scaffolded, lysosomal p62 oligomers are highly stable.

**Figure 5.**
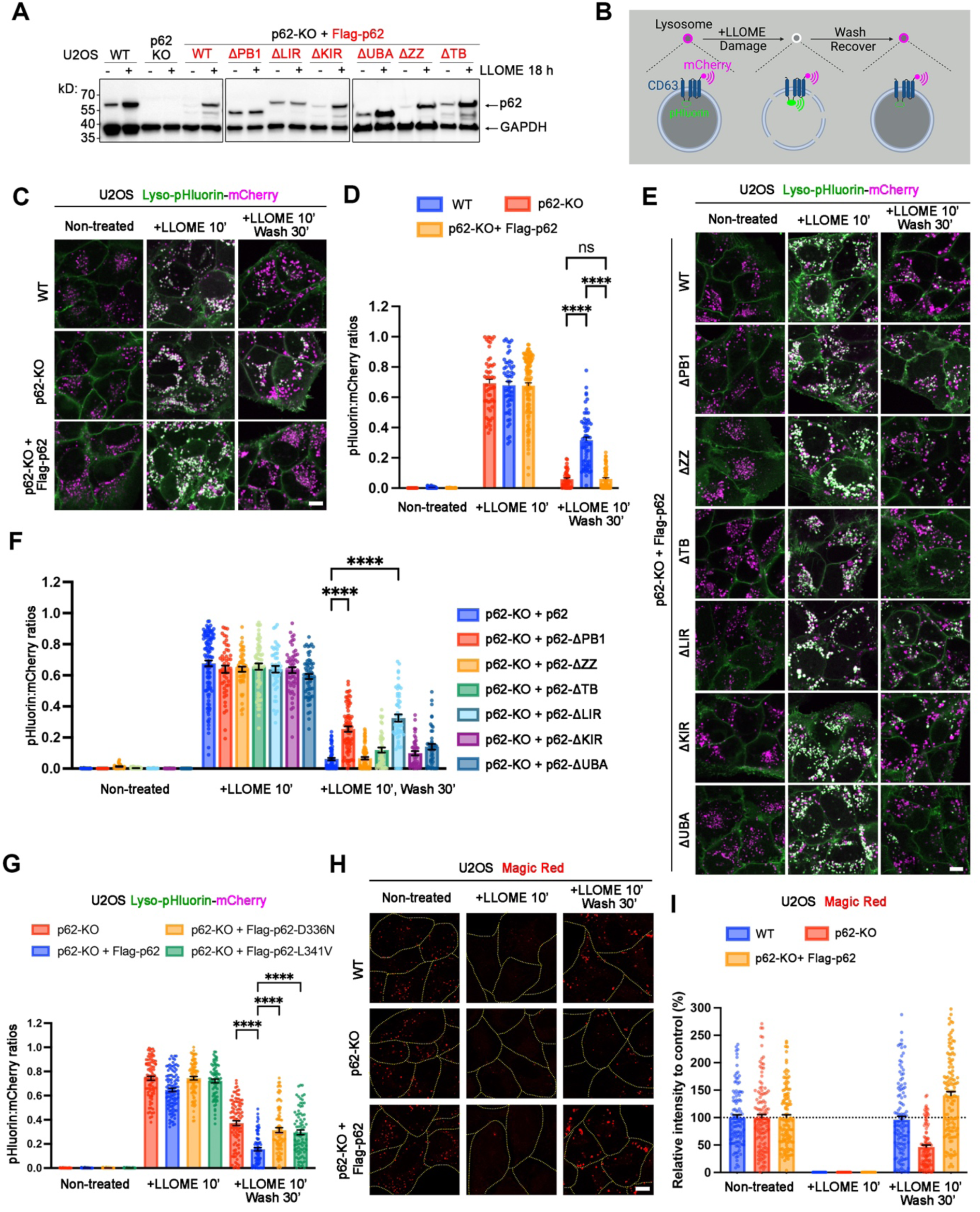
p62 promotes lysosomal repair. (A) Immunoblot showing that LLOME increases the protein levels of endogenous and re-expressed p62 in U2OS cells. (B) Schematic illustration of lysosomal repair assay using cells stably expressing lyso-pHluorin-mCherry. The pH-sensitive GFP variant pHluorin (green) was inserted into a luminal loop of CD63 and its signal is quenched by the acidic lysosomal pH when the lysosome is intact or repaired; mCherry (red) was linked to the C-terminal end of CD63. (C) Confocal images of the lysosomal repair assay showing repair defects of p62-KO cells, which are rescued by p62 re-expression. (D) Quantification of the pHluorin-to-mCherry signal ratios in (C). Data are means ± SEM; n = 18, 42, 60, 51, 54, 110, 43, 62, 72 random cells, respectively. (E) Confocal images showing that the lysosomal repair defects of p62-KO cells are rescued by the re-expression of p62 and its mutants except for ΔPB1 and ΔLIR, the self-oligomerization and ATG8-binding mutants, respectively. (F) Quantification of the pHluorin-to-mCherry signal ratios in (E). Data are means ± SEM; n = 60, 27, 36, 79, 30, 34, 30, 110, 49, 52, 62, 46, 52, 62, 72, 72, 81, 46, 54, 45, 45 random cells, respectively. (G) Quantification of the pHluorin-to-mCherry signal ratios in S6(E). Data are means ± SEM; n = 66, 51, 53, 70, 90, 105, 93, 76, 96, 76, 86, 84 random cells, respectively. (H) Magic Red assay showing that p62-KO inhibits the recovery of lysosomal cathepsin activity. Cells were treated as indicated and then Magic Red (1:250) was added, followed by live cell imaging. (I) Quantification of the relative Magic Red signals in (G). Data are means ± SEM; n = 107, 123, 116, 94, 104, 100, 119, 104, 126 random cells, respectively. Data were analyzed by two-way ANOVA with Tukey’s multiple comparison test for (D), (F), (H) and (I). WT, wild type. KO, knockout. Bar, 10 μm. See also Figure S6.

The LMP-induced p62 stabilization suggests potential roles of p62 oligomers in lysosomal membrane stabilization. In addition, protein oligomerization, such as ESCRT-III assemblies and annexins, are known to mediate lysosomal membrane repair. Furthermore, the CASM-dependent assembly of both p62 and ESCRT oligomers suggests that p62 oligomers might play a role in lysosomal repair similarly to ESCRT.

To quantitatively analyze the impact of p62 on lysosomal repair, we employed a genetically encoded ratiometric lysosomal pH sensor, Lyso-pHluorin-mCherry^55^ (**Figure 5B**). In resting conditions, with the quenching of pHluorin signal by the acidic lysosomal pH, the sensor only showed red puncta (**Figure 5B, C**). Following LLOME-mediated lysosomal damage, pHluorin was activated and colocalized with the mCherry puncta, which was then reversed as lysosomes were repaired following LLOME wash out (**Figure 5B, C**). A substantial delay in lysosomal repair was captured in p62-KO cells, which was fully rescued by p62 reconstitution in the KO cells (**Figure 5C, D**). Consistent with a key role for CASM in recruiting both ESCRT and p62, knockout of either *ATG5* or *ATG7* caused a dramatic defect of rapid lysosomal repair, as assessed by the ratiometric lysosomal pH sensor (**Figure S6A** and **S6B**).

To further assess whether the lysosomal recruitment of p62 is essential for its role in lysosomal repair, we reconstituted each of the p62 deletion mutants (**Figure 3F**) in p62-KO cells to examine their impact on lysosomal repair. All the p62 mutants rescued lysosomal repair except ΔPB1 and ΔLIR (**Figure 5E, F**), the two mutants that were no longer recruited to damaged lysosomes (**Figure 3G** and **H**). These data strongly support that p62 oligomerization facilitates lysosomal membrane repair.

Loss-of-function mutations of p62 are commonly found in multiple human diseases, such as metabolic disease, skeletal and muscular disorders, and neurodegenerative diseases, particularly amyotrophic lateral sclerosis (ALS) & frontotemporal dementia (FTD)^56^. Two ALS/FTD-associated p62 mutations, L341V and D336N, largely lost LC3 interaction in co-immunoprecipitation^57^. Consistent with a key role for ATG8 interaction in p62 recruitment, the two disease-associated point mutations each strongly suppressed LMP-induced lysosomal recruitment of p62 (**Figure S6C** and **D**). Both mutations also failed to rescue lysosomal repair in p62-knockout cells, as shown by the ratiometric lyso-pHluorin-mCherry probe (**Figure S6E** and **5G**). Thus, A loss-of-function of p62 in lysosomal repair is likely underlying ALS and FTD.

We further asked whether p62-mediated lysosomal repair correlated to the restoration of lysosomal hydrolytic activity. Using Magic Red, a fluorogenic peptide substrate that becomes fluorescent only after cleavage by active lysosomal cathepsins^58^, we were able to analyze lysosomal protease activity under live cell imaging. Lysosomal damage by LLOME completely eliminated Magic Red fluorescence signals, which fully recovered after LLOME washout (**Figure 5H, I**). Knockout of p62 substantially reduced the recovery of Magic Red signals, which was fully rescued by the reconstitution of p62 in the knockout cells (**Figure 5H, I**). Together, these results are consistent with a role of p62 in lysosomal membrane repair, likely achieved through membrane stabilization by CASM-scaffolded p62 oligomerization.

### The CASM-p62 axis recruits KEAP1 to damaged lysosomes

To explore additional functions of p62 on damaged lysosomes beyond membrane repair, we analyzed p62 interactome by TurboID-mediated proximity labeling (**Figure 6A**). p62-specific interacting proteins were identified by comparing the hits of TurboID-p62 and Lyso-TurboID^14^ following lysosomal damage by LLOME (**Figure 6A**). Among the very top hits were p62 itself, tubulin alpha 1C and beta 4B (TUBA1C, TUBB4B), ubiquitin (RPS27A), and Kelch-like ECH-associated protein 1 (KEAP1), a cytosolic redox sensor (**Figure 6B, Supplementary Table S3**). While lysosome damage has been known as a trigger for cellular oxidative stress^59^, how cells handle such stress for detoxication remains unclear. We therefore explored whether p62 recruits KEAP1 and thereby regulates redox signaling in response to lysosomal damage.

**Figure 6.**
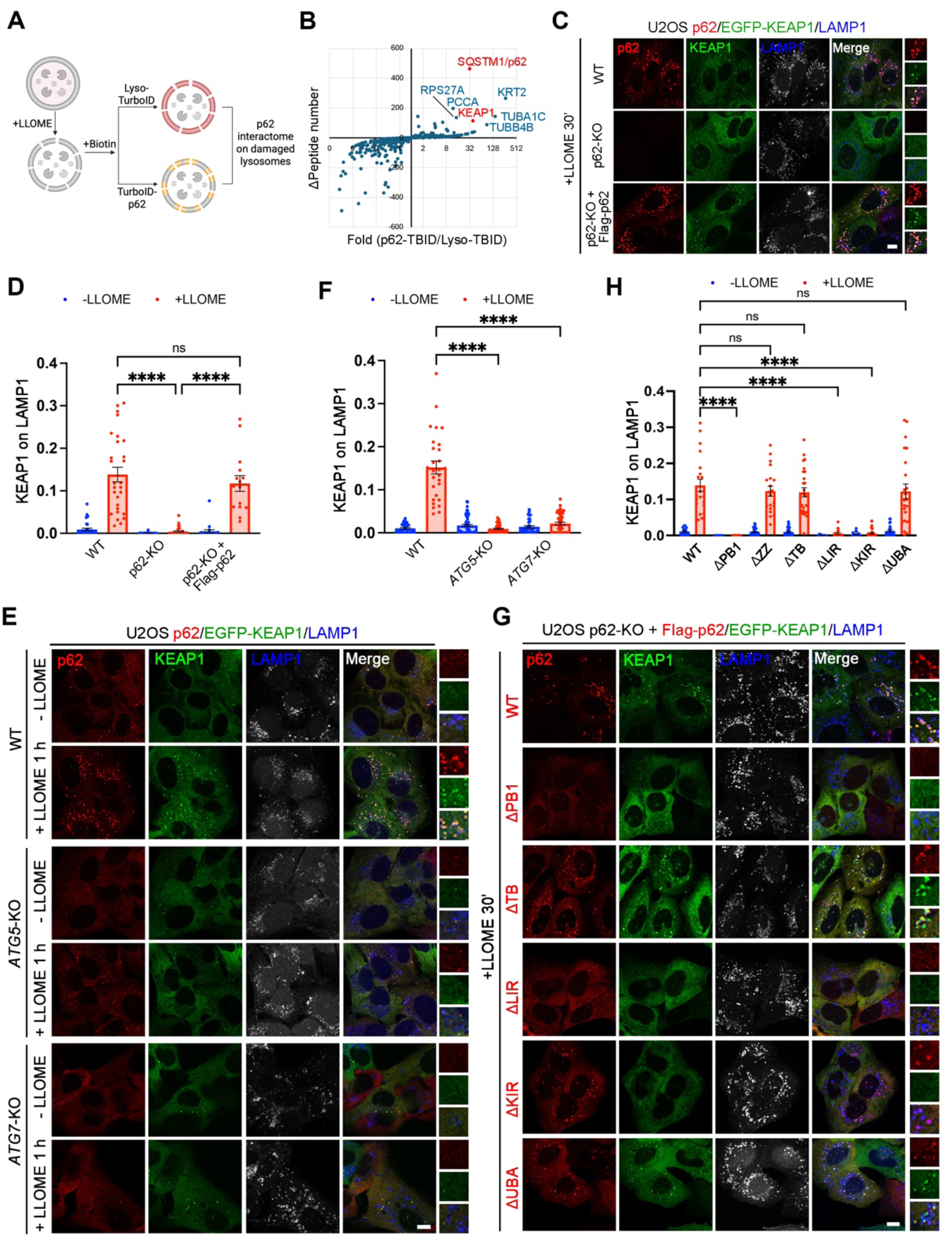
The CASM-p62 axis recruits KEAP1 to damaged lysosomes. (A) Schematic illustration of the TurboID screen searching for p62-interacting proteins on damaged lysosomes. (B) Dot plot of proteins identified from Lyso-TurboID and p62-TurboID following lysosomal damage by LLOME. (C) p62 mediates KEAP1 recruitment to damaged lysosomes. U2OS cells stably expressing EGFP-KEAP1 were treated with 500 μM of LLOME for 30 min, followed by fixation and immunostaining of endogenous LAMP1 and p62. (D) Quantification of EGFP-KEAP1 recruitment to lysosomes in (C). Data are means ± SEM; n = 31, 28, 39, 33, 22 15random cells, respectively. (E) Knockout of ATG5 or ATG7 blocks the recruitment of p62 and EGFP-KEAP1 to damaged lysosomes. Indicated U2OS cells stably expressing EGFP-KEAP1 were treated with or without 500 μM of LLOME for 1 h, followed by fixation and immunostaining of endogenous LAMP1 and p62. (F) Quantification of EGFP-KEAP1 recruitment to lysosomes in (E). Data are means ± SEM; n = 30, 28, 62, 47, 36, 48 random cells, respectively. (G) Deletion of the KIR domain of p62 blocks LLOME-induced lysosomal recruitment of KEAP1. U2OS p62-KO cells, stably expressing EGFP-KEAP1 together with p62 or its mutants, were treated with 500 μM of LLOME for 30 min, followed by fixation and immunostaining of Flag-p62 and endogenous LAMP1 62. (H) Quantification of EGFP-KEAP1 recruitment to lysosomes in (G). Data are means ± SEM; n = 16, 21, 23, 21, 36, 20, 28, 26, 12, 17, 22, 37, 24, 23 random cells, respectively. Data were analyzed by two-way ANOVA with Tukey’s multiple comparison test for (D), (F) and (H). WT, wild type. KO, knockout. Bar, 10 μm. See also Figure S7.

Stably expressed EGFP-KEAP1 showed even cytosolic distribution in U2OS cells at rest (**Figure S7A**). Lysosomal damage triggered the formation of KEAP1 puncta that strongly colocalized with p62 lysosomes (**Figure 6C, D**). Knockout of p62 completely abolished LMP-induced KEAP1 puncta, which was restored by p62 reconstitution in the knockout cells (**Figure 6C, D**).

Consistent with a key role for lysosomal CASM in p62 recruitment, *ATG5*-KO and *ATG7*-KO cells lost KEAP1 recruitment to damaged lysosomes (**Figure 6E, F**). Compound TAK243, which almost completely eliminated cellular ubiquitination (**Figure S1L**) and abolishes lysosomal recruitment of other autophagy adaptors (**Figure S2C-F**), did not affect LMP-induced KEAP1 recruitment (**Figure S7B, C**). Thus, KEAP1 is recruited to damaged lysosomes through a p62-dependent, ubiquitin-independent pathway downstream of lysosomal CASM.

To further illustrate how p62 recruits KEAP1 to damaged lysosomes, we reconstituted various domain deletion mutants of p62 in the knockout cells. p62 carries a KEAP-interacting region (KIR) in its longest IDR which was previously shown to directly bind KEAP1^60,61^. Deletion of KIR (ΔKIR) did not affect p62 targeting to damaged lysosomes, but fully abrogated KEAP1 recruitment (**Figure 6G, H; S7D**). Besides the KIR domain, deletion of the PB1 or LIR domain that affected p62 recruitment also eliminated KEAP1 recruitment to damaged lysosomes (**Figure 6G, H; S7D**). These findings establish KEAP1 as a direct effector of the lysosomal CASM-p62 axis during lysosome damage.

### The lysosomal CASM-p62-KEAP1 axis activates NRF2

NRF2 (NFE2L2) is a stress-inducible transcription factor that activates antioxidant and cytoprotective gene expression^62^. NRF2 is negatively regulated by KEAP1, a substrate adaptor for a Cul3-based E3 ubiquitin ligase complex that promotes NRF2 ubiquitination and proteasomal degradation^63–67^. Furthermore, p62 competes with NRF2 for KEAP1 binding, which in turn activates NRF2 (ref^61^). We explored whether the lysosomal CASM-p62-KEAP1 axis regulates NRF2 activation. Indeed, lysosomal damage by LLOME enhanced the nuclear localization of endogenous NRF2 (**Figure 7A, B, S7E**), an indication of NRF2 activation^63^. Knockout of p62 substantially suppressed LMP-induced lysosomal NRF2 translocation, which was fully rescued by the reconstitution of wild-type p62 in the knockout cells (**Figure 7A, B, S7E**). Thus, p62 appears to play a key role in LMP-induced NRF2 activation.

**Figure 7.**
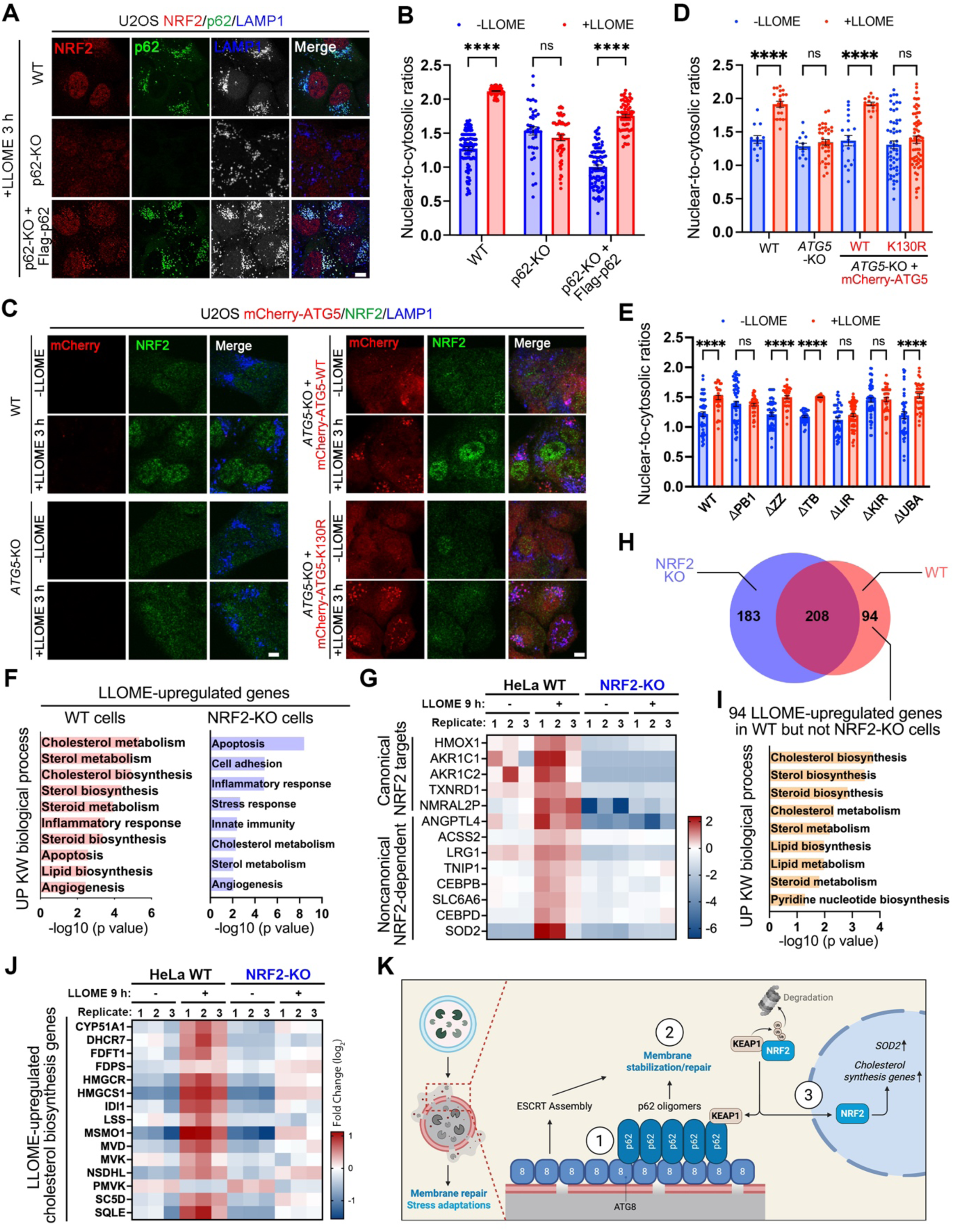
SOLAR activates NRF2-dependent redox defense and cholesterol synthesis. (A) LLOME-induced NRF2 nuclear translocation is p62 dependent. U2OS WT and p62-KO/rescue cells were treated with 500 μM of LLOME for 3 h, followed by fixation and immunostaining of endogenous NRF2, p62 and LAMP1. (B) Quantification of NRF2 nuclear translocation in (A). Data are means ± SEM; n = 106, 75, 40, 53, 87, 64 random cells, respectively. (C) LLOME-induced NRF2 nuclear translocation is dependent on ATG5 and its conjugation to ATG12. Indicated U2OS cells were treated with 500 μM of LLOME for 3 h, followed by fixation and immunostaining of endogenous NRF2 and LAMP1. ATG5 K130R is deficient in ATG12 conjugation. (D) Quantification of NRF2 nuclear translocation in (C). Data are means ± SEM; n = 13, 21, 14, 35, 20, 14, 56, 60 random cells, respectively. (E) Quantification of NRF2 nuclear translocation in Fig S7F. Data are means ± SEM; n = 44, 45, 61, 39, 49, 38, 34, 29, 46, 63, 53, 35, 40, 39 random cells, respectively. (F) UniProt (UP) keyword (KW) biological process analysis of LLOME-upregulated genes in WT and *NRF2*-KO HeLa cells. (G) Heatmap displaying the expression pattern of canonical and noncanonical NRF2-upregulated genes in WT and *NRF2*-KO HeLa cells. (H) Venn diagram showing the number of genes depending on NRF2 for LMP-induced upregulation (log2FC >=1, p value < 0.01). (I) UP KW biological process analysis of LLOME-upregulated genes in WT but not NRF2-KO HeLa cells. Note that “cholesterol biosynthesis” is the only strongly enriched pathway. (J) Heatmap displaying the expression pattern of genes from the cholesterol biosynthesis pathway in WT and *NRF2*-KO HeLa cells. (K) Schematic illustration of the SOLAR pathway. Lysosomal membrane damage activates the conjugation of ATG8s to lysosomal membranes (CASM), which in turn recruits p62 through ATG8-p62 binding and p62 self-oligomerization. The assembly of p62 through CASM contributes to lysosomal membrane repair, in parallel with the ESCRT machineries. The CASM-p62 axis also activates NRF2 through KEAP1 recruitment, leading to NRF2-dependent upregulation of mitochondrial antioxidative enzyme SOD2 and cholesterol biosynthesis genes. Data were analyzed by two-way ANOVA with Tukey’s multiple comparison test for (B), (D), and (E). WT, wild type. KO, knockout. Bar, 10 μm. See also Figure S7 and S8.

In support of CASM as a driver of p62-dependent NRF2 activation in response to lysosomal damage, we found that *ATG5* knockout abolished LMP-induced nuclear translocation of NRF2 (**Figure 7C, D**). Reconstitution of wild-type but not the conjugation-deficient mutant of ATG5 fully restored LMP-induced NRF2 nuclear translocation (**Figure 7C, D**). Note that both wild-type and mutant ATG5 were recruited to damaged lysosomes following LLOME treatment (**Figure 7C**), consistent with CASM activation.

To examine whether p62 activated NRF2 through the CASM-p62-KEAP1 axis, we re-expressed various p62 mutants in the p62-KO cells. The two p62 mutants deficient in lysosomal recruitment, ΔPB1 and ΔLIR, failed to rescue LMP-induced NRF2 activation (**Figure 7E and S7F**). Similarly, ΔKIR was not able to support NRF2 activation following lysosomal damage, whereas the remaining mutants all restored NRF2 activation (**Figure 7E and S7F**). Thus, p62 recruitment through CASM and its binding to KEAP1 contributes to NRF2 activation following lysosomal damage.

Our findings reveal CASM-dependent lysosomal p62 recruitment as a mechanistically robust and functionally unique pathway on damaged lysosomes, different from the canonical ubiquitination-dependent recruitment of autophagy adaptors. To distinguish this pathway from established roles of p62 and related adaptors in autophagy, we designate it SOLAR (SQSTM1/p62 oligomer-mediated lysosomal antioxidant defense and membrane repair). The SOLAR pathway represents a CASM-dependent lysosomal defense program in which p62 oligomers coordinate membrane repair while concurrently promoting NRF2-mediated antioxidant signaling. This dual function differentiates SOLAR from ESCRT-mediated membrane repair and ubiquitination-driven lysophagy.

### LMP activates NRF2-dependent redox defense, metabolic adaptation and cholesterol biosynthesis

LMP is known to trigger oxidative stress through multiple mechanisms, including the release of redox active iron and cathepsin-mediated mitochondrial dysfunction^59,68^. Accordingly, SOLAR-dependent activation of NRF2, an established antioxidative transcription factor^62^, is expected to promote redox defense. To analyze the LMP-induced transcriptional function of NRF2, we performed RNA-seq experiments using wild-type and *NRF2*-KO HeLa cells. LMP induced by LLOME treatment triggered the transcriptional upregulation of over 300 genes (**Figure S8A-D**), enriched in genes implicated in inflammatory response and apoptosis pathways (**Figure 7F**), consistent with previous reports^69^. Most of the previously established NRF2 targets such as *SLC7A11, GCLC, GCLM, NQO1,* and *SOD1* appeared to have high basal expression in HeLa cells in a NRF2-dependent manner and were not further upregulated by LLOMe treatment (**Figure S8E**). By contrast, a group of highly-inducible NRF2 targets were strongly upregulated by LLOMe in a NRF2-dependent manner, including *HMOX1, AKR1C1, AKR1C2, TXNRD1,* and *NMRAL2P* (**Figure 7G**), all of which are well-established NRF2-responsive genes involved in antioxidant defense and electrophile detoxification^70–72^. In addition to canonical redox defense genes, lysosomal membrane damage by LLOMe also upregulated a list of noncanonical targets in a NRF2-dependent manner, which are associated with metabolic adaptation and mitochondrial stress resistance, including *ANGPTL4, ACSS2, LRG1, TNIP1, CEBPB, SLC6A6, CEBPD,* and *SOD2* (**Figure 7G**). Among them, mitochondrial superoxide dismutase 2 (*SOD2*), a more recently identified NRF2-upregulated gene^73–75^, is markedly upregulated by LLOMe, which was observed in both RNA-seq and quantitative PCR (q-PCR) (**Figure S8F**). Thus, lysosomal damage triggers a selective NRF2-dependent adaptive program rather than a broad canonical antioxidant response.

To further examine the transcriptional function of NRF2 in response to LMP, we performed pathway enrichment analysis using the total list of 94 genes upregulation by LLOME in a NRF2-dependent manner (**Figure 7H, Supplementary Table S4**). Cholesterol biosynthesis was identified as the most significantly enriched pathway among these genes (**Figure 7I**). Further analysis revealed that 14 out of the 15 genes in the cholesterol biosynthesis pathway were upregulated by LLOME (**Figure S8G**), and such upregulation was largely gone in *NRF2*-KO cells (**Figure 7J**). Given the critical role of ER-to-lysosome cholesterol transfer in lysosomal membrane repair^13,14^, increased cholesterol synthesis likely replenishes ER cholesterol and enhances cellular capacity for membrane repair.

Together, lysosomal membrane damage triggers a NRF2-dependent transcriptional program characterized by the upregulation of selective NRF2-targets for redox defense, metabolic adaptation, and cholesterol biosynthesis.

## DISCUSSION

Lysosomal damage is emerging as a common driver of aging pathology^1^. Illustrating the molecular architecture of lysosomal damage response is crucial for future therapeutic interventions targeting lysosomes. In this study, we define SOLAR as a CASM-dependent lysosomal quality control pathway that links membrane repair with cholesterol synthesis and redox adaptation (**Figure 7K**). LMP-induced Atg8ylation serves as a platform to recruit and stabilize p62 oligomers on damaged lysosomes, through coordinated ATG8 binding and p62 self-assembly. This recruitment is independent of ubiquitination at early stages and is required for efficient lysosomal membrane repair. In parallel, p62 recruits KEAP1 and activates NRF2, turning on a transcriptional response upregulating cholesterol biosynthesis and cellular redox defense. Given the additional role of p62 in lysophagy^7,34^, it seems likely that the SOLAR pathway may participate in determining the fate of damaged lysosomes. We speculate that early CASM-dependent recruitment of p62 may favor membrane repair and recovery, whereas later ubiquitin-dependent recruitment may shift p62 function toward lysophagy, likely involving p62 condensation^34,76^.

A major question in the field of lysosomal quality control is how the diverse repair pathways are coordinated. CASM is emerging as a platform for multiple lysosomal repair pathways^1^, including the SOLAR pathway described here. Both ESCRT^52–54^ and SOLAR are recruited to damaged lysosomes through a CASM-dependent manner. The dramatically enhanced recruitment of p62 in ESCRT-depleted cells suggests that ESCRT and SOLAR may play complementary roles in lysosomal repair. CASM also cross-talks with other repair machineries such as ATG2 (ref^43^) and BLTP3A^77^, both bridge-like lipid transfer proteins implicated in lysosomal repair through lipid transfer. Of note, primary neurons appear to be deficient in ESCRT assembly in response to lysosomal membrane damage^78^, which may be partially explained by reduced CASM activity in neurons^79^. Our findings suggest that such ESCRT assembly defect may drive strong p62 recruitment for lysosomal membrane repair in neurons.

Given that both lysosomal dysfunction and oxidative stress are central features of aging and age-related diseases^80^, the SOLAR pathway may have important implications in aging biology. Indeed, p62 levels are reduced in multiple aged tissues^81,82^, and knockout of p62 accelerates aging phenotypes^83^. The lysosomal repair defects caused by two ALS/FTD-related p62 mutations suggest a potential link between impaired SOLAR assembly and neurodegeneration. Furthermore, loss of p62 increases mitochondrial ROS production and impairs respiratory function^83^, consistent with a role for SOLAR in upregulating the mitochondrial antioxidative enzyme SOD2. Notably, while acute SOALR activation is expected to promote lysosomal repair and redox homeostasis, the impact of chronic lysosomal dysfunction on NRF2-dependent metabolic reprogramming and cholesterol synthesis in aging and disease contexts remains to be fully defined. In summary, our findings reveal SOLAR as an LMP-induced platform coordinating lysosomal membrane repair, cellular redox defense, and de novo cholesterol biosynthesis, with potential implications for aging and neurodegenerative disease biology.

## RESOURCE AVAILABILITY

Requests for further information and resources should be directed to and will be fulfilled by the lead contact, Jay Xiaojun Tan.

### Materials availability

Plasmids generated in this study will be deposited to Addgene and will be available upon the publication of the manuscript. All other unique/stable reagents generated in this study are available from the lead contact with a completed materials transfer agreement.

### Data and code availability

The mass spectrometry data have been deposited to the ProteomeX-change Consortium via the PRIDE partner repository and will be publicly available as of the date of publication. RNA seq raw data are available through NCBI GEO (GSE329018). All other data are provided within the paper and its Supplementary Information. Original western blot images will be deposited at Mendeley and will be publicly available as of the date of publication. All customized code for image quantification has been deposited to an open-source repository at: https://github.com/jaytanlab/Protein_Colocalization_Quantification. Any additional information is available from the lead contact upon request.

## ACKNOWLEDGMENTS

We thank members of the Tan lab and the Aging Institute at the University of Pittsburgh for discussions; MS Bioworks for mass spectrometry services. Schematic illustrations were generated using Biorender. This work was supported by start-up funding from the Aging Institute at the University of Pittsburgh School of Medicine and University of Pittsburgh Medical Center (UPMC) to J.X.T.

## AUTHOR CONTRIBUTIONS

Conceptualization, J.X.T.; methodology, J.X., Z.Y., H.Y., J.C., X.Y., J.X.T.; Investigation, all authors; writing—original draft, J.X.T. with input from co-authors; writing—review & editing, all authors; funding acquisition, J.X.T.; supervision, J.X.T.

## SUPPLEMENTAL INFORMATION

Supplementary Table S1. ATG8-interacting proteins enriched on damaged lysosomes. Related to Figure 1A. Supplementary Table S2. DNA oligo sequences.

Supplementary Table S3. p62-interacting proteins enriched on damaged lysosomes, Related to Figure 6B. Supplementary Table S4. LLOME upregulated genes in WT and *NRF2*-KO cells. Related to Figure 7H.

## Materials and Methods

### Cell culture, chemicals, and treatments

U2OS, HeLa, 293T, HT1080, BEAS-2B, and PC3 cells were originally from ATCC and were authenticated through short tandem repeat profiling and their profiling data are publicly available from ATCC. The morphologies of these cell lines differ from each other, and their growth rates and morphologies were routinely monitored. All cells were tested by PCR to ensure they were free of mycoplasma contamination. U2OS, HeLa, 293T, HT1080, BEAS-2B cells were maintained in Dulbecco’s modified Eagle’s medium (DMEM); PC3 cells were cultured in Roswell Park Memorial Institute (RPMI) 1640. The media were supplemented with 10% FBS, 100 U/ml Penicillin-Streptomycin, and 2.5 mg/ml plasmocin, at 37°C with 5% CO_2_.

LLOME (Sigma, #L7393), Monensin (Cayman Chemical, 16488), and nigericin (Cayman Chemical, #11437) were dissolved in ethanol and stored in aliquots at -20°C. Torin1 (Cayman Chemical, #10997), TAK243 (MedChemExpress, #HY-100487), HOIPIN-8 (Selleckchem #E1798), NSC697923 (Selleckchem, #S7142), Rapamycin (Cayman Chemical, #13346) and ML-SA1 (Cayman Chemical, #29958) were dissolved in DMSO and stored at -20°C. For cell treatments, the compound stocks were directly diluted into the cell culture media. When media change is needed, media prewarmed in the same incubator were used. Magic Red was directly added to the culture media.

### Antibodies

Rabbit anti-OSBP (11096-1-AP, immunoblot 1:1,000, immunofluorescence 1:1,000), rabbit anti-ORP9 (11879-1-AP, immunoblot 1:1,000), rabbit anti-LC3 (14600-1-AP, immunoblot 1:1,000, immunofluorescence 1:1,000), rabbit anti-GABARAP (18723-1-AP, immunoblot 1:1,000), rabbit anti-ATG7 (81760-1-RR, immunoblot 1:1,000), rabbit anti-IST1 (19842-1-AP, immunofluorescence 1:2,000), rabbit anti-Galectin3 (14979-1-AP, immunofluorescence 1:1,000), and mouse anti-GAPDH (60004-1-Ig, immunoblot 1:1,000) were from Proteintech. Mouse anti-ORP9 (A-7, sc-398961, immunofluorescence 1:1,000), mouse anti-PI4K2A (sc-390026, immunofluorescence 1:1,000), mouse anti-LAMP2 (sc-18822, immunofluorescence 1:1,000), mouse anti-LAMP1 (sc-20011, immunofluorescence 1:1,000), mouse anti-p62(sc-28359, immunoblot 1:1,000, immunofluorescence 1:1,000), Mouse Anti-Ubiquitin (sc-271289, immunofluorescence 1:1,000) were from Santa Cruz Biotechnology. Rabbit anti-LAMP1 monoclonal antibody (9091S, immunofluorescence 1:1,000), Rabbit anti-NRF2 monoclonal antibody (12721S, immunofluorescence 1:1,000), Rabbit anti-Beta-Actin (4970S, immunoblot 1:1,000), Rabbit anti-ATG13 (13468, immunoblot 1:1,000), Rabbit anti-ATG16L1 (8089S, immunoblot 1:1,000) were from Cell Signaling. Flag (M2, immunofluorescence 1:10,000) and Rabbit anti-ATG5 (A0731, immunoblot 1:1,000) were from Sigma. Mouse anti-CD63 (556019, immunofluorescence 1:1,000) was from BD Pharmingen. Guinea anti-p62 (GP62-C, immunofluorescence 1:1,000) was from Progen. Mouse Anti-ALIX (634502, immunofluorescence 1:1,000) was from Biolegend. Alexa-488/594- and Pacific Blue-conjugated secondary antibodies were from ThermoFisher Scientific.

### DNA cloning

Stable protein expression was achieved by a lentiviral approach. The relevant DNA sequences were cloned into pCDH-CMV-MCS and the peptide linker GSGSGS was used when fusing two sequences together. When introducing point mutations, PCR was applied to generate DNA fragments which carried the desired mutations at their overlapped ends and then the fragments were fused together into pCDH vector through infusion. All cDNA sequences were of human origin unless otherwise specified. KEAP1 cDNA was amplified from pDsRed2-Keap1 (Addgene, #21551)^1^. ATG5 cDNA was amplified from mCherry-ATG5-C-18 (Addgene, # 54995). ATG16L1 cDNA was amplified from a Myc-ATG16L1 plasmid shared by Qing Zhong. SQSTM1/p62 cDNA was amplified from pET-Duet1_6xHis-TEV-eGFP-P62 (Addgene, #190929)^2^. pCDH-EGFP-ATG2A plasmid was previously described^3^. Primer sequences for DNA cloning are summarized in **Table S2**.

### Stable cell line generation

All experiments were based on stable cell lines with genetic knockouts and/or stable ectopic protein expression through a lentiviral approach, except siRNA-mediated knockdown which was based on transient transfection. pCDH vectors were used to generate lentiviruses carrying specific DNA sequences of interest to infect recipient cells for stable expression. Puromycin was used to eliminate uninfected cells when needed, but this was often unnecessary given the nearly 100% infection rates. The process of viral titration was carried out to identify the smallest virus titers required for inducing target protein expression in over 90% of cells.

To generate knockout cell lines, Cas9 guide sequences were incorporated into LentiCRISPR.v2 (Addgene #52961) for lentivirus packaging (*67*). A variety of guide sequences were evaluated for each gene to identify the most effective one that significantly diminished target expression in the CRISPR knockout pools. Primers containing guide sequences for CRISPR cloning are summarized in **Table S2**.

### Biotinylation and purification of lysosomal surface proteins

Lysosomal surface biotinylation using Lyso-TurboID was previously described^3^. To identify p62 interactome on lysosomal surface following lysosomal membrane damage, 293T cells with stable expression of p62-TurboID were cultured to 80% confluence. Subsequently, the cells were exposed to 500 μM LLOME or a vehicle for 30 min to induce the lysosomal membrane permeabilization. After this treatment, 50 μM biotin was introduced and allowed to incubate for 30 min to biotinylate proteins on the lysosomal surface. The cells were subsequently washed twice with pre-chilled PBS, scraped into 1 ml of cold PBS per dish, and moved into 1.5-ml tubes. After being resuspended in 1 ml of hypotonic buffer (10 mM HEPES, pH 7.5, 5 mM MgCl2, 10 mM NaCl), the cells were passed through a 25 G needle ten times and centrifuged at 1,000g for 3 min to remove intact cells and nuclei. The supernatant (S1) underwent additional centrifugation at 20,000g for 20 min to precipitate the majority of membranes, including lysosomes with biotinylated proteins (P20). To completely dissolve the membranes, P20 was resuspended in 500 ul lysis buffer (50 mM Tris-HCl, pH 8.0, 150 mM NaCl, 0.5% Triton X-100, 2 mM NaF, 5 mM MgCl2, protease inhibitor cocktail) and briefly sonicated. Following an additional centrifugation at 15,000g for 5 min to eliminate undissolved aggregates, the supernatants were transferred to a new tube containing 100 μl of well-resuspended streptavidin magnetic beads and rotated at 4°C for 2 h. The beads were then washed twice with each of the following three buffers sequentially: buffer 1 (50 mM Tris-HCl, pH 8.0, 150 mM NaCl, 0.8% SDS), buffer 2 (50 mM Tris-HCl, pH 8.0, 1 M NaCl, 0.5% Triton X-100), and buffer 3 (50 mM Tris-HCl, pH 8.0, 150 mM NaCl, 0.5% Triton X-100). Proteins bound to the beads were eluted by adding 50 μl of 2×SDS sample buffer (0.1 M Tris-HCl, pH 6.8, 4% SDS, 20% glycerol, 2% 2-mercaptoethanol, 0.01% bromphenol blue), heated at 95°C for 5 min, and stored at -80°C.

### Immunofluorescence

After being seeded onto glass coverslips (Warner Instruments, 64-0712) in 24-well plates for 24 to 48 h, cells were treated as indicated. After treatment, cells were fixed using 4% paraformaldehyde (PFA) in PBS for 30 min at room temperature, followed by permeabilization with 0.1% Triton X-100 for 2 min. Afterwards, they were incubated with a 1x blocking buffer (ThermoFisher Scientific, 37565) for 1 h at room temperature. The same buffer was used to dilute both primary and secondary antibodies. The cells were then incubated with primary antibodies diluted in the same 1x blocking buffer overnight at 4°C. After three washes with PBS, the cells were incubated with secondary antibodies for 30 to 60 min at room temperature, followed by three additional washes with PBS. A quick 2-min DAPI stain was applied as necessary. Finally, coverslips were mounted onto slides using VECTASHIELD HardSet Antifade Mounting Medium (Vector Laboratories, H-1700) and left to stabilize for more than an hour at room temperature before imaging. The slides were stored at -20°C.

Images were captured using a Leica SP8 LIGHTNING confocal system, utilizing the built-in Leica Application Suite X 3.5.5.19976 software. Negative controls were employed to rule out nonspecific staining and channel crosstalk. Live-cell imaging on the same confocal system was conducted using an Okolab stage-top incubator, which maintained consistent temperature and CO_2_ concentration. The same software settings and Adobe Photoshop were used for all image processing within the same figure panel. Custom software codes automatically analyzed images in a blinded fashion.

### Immunoblotting

Cells were washed with pre-chilled PBS and then lysed in a buffer containing 50 mM Tris-HCl (pH 7.5), 150 mM NaCl, 0.5% Triton X-100, 2 mM NaF, 5 mM MgCl₂, and a protease inhibitor cocktail. The lysates were briefly sonicated and centrifuged at 15,000g for 10 min. The resulting supernatants were collected and mixed with an equal volume of 2×SDS loading buffer (0.1 M Tris-HCl, pH 6.8, 4% SDS, 20% glycerol, 2% 2-mercaptoethanol, 0.01% bromophenol blue), then heated at 95°C for 10 min. Equal amounts of total proteins from each sample were separated by sodium dodecyl sulfate–polyacrylamide gel electrophoresis (SDS–PAGE) using Mini-PROTEAN TGX Precast Gels (4–20%, Bio-Rad, # 4561091). The proteins were then transferred onto 0.22 μm Trans-Blot Turbo nitrocellulose membranes. The membranes were blocked with StartingBlock Blocking Buffer (ThermoFisher, #37542) for 1 h at room temperature, followed by overnight incubation at 4°C with primary antibodies diluted in the same blocking buffer. After washing with PBS containing 0.1% tween, the membranes were incubated for 1 h at room temperature with HRP-conjugated secondary antibodies. Protein signals were developed using Immobilon Forte chemiluminescent substrates (Millipore, #WBLUF0100) and visualized using the Invitrogen iBright FL1000 Imaging System (ThermoFisher Scientific).

### Lysosomal repair assay

Lysosomal repair was examined in cells stably expressing Lyso-pHluorin-mCherry, a ratiometric probe for lysosomal pH^4^. This is a CD63-based probe carrying a pH-sensitive pHluorin in a luminal loop and mCherry at its C-terminal end on the lysosomal surface. In resting conditions, the pHluorin signal is quenched by acidic pH of lysosomes, showing red-only puncta on lysosomes. Lysosomal damage activates pHluorin signals, which colocalize with mCherry and appear as yellow puncta that return to red upon lysosomal repair and reacidification. Cells were seeded in 24-well plates, and 48 h later cells were treated with xx mM LLOME for 10 min, followed by media change (Wash). Lyso-pHluorin-mCherry signals were captured by live cell confocal imaging.

### RNA isolation for PCR and RNA sequencing

Cells were harvested upon their respective treatments and washed twice with cold 1× PBS and subsequently scraped into RNase-free tubes. The samples were centrifuged at 1000 × g for 3 minutes at 4 °C to pellet the cells, and the supernatant was carefully aspirated. The resulting cell pellets were resuspended in 600 µL of 1% 2-mercaptoethanol (2-ME) lysis buffer (Purelink RNAmini, Invitrogen, cat #12183018) and homogenized by passing each sample through a 21-gauge needle eight times. An equal volume (600 µL) of 70% ethanol was then added, and the mixtures were transferred to spin columns. Columns were washed by centrifugation at 12,000 × g for 15 seconds following the addition of wash buffer. DNase treatment buffer (Purelink^TM^ DNase, Invitrogen, cat #12185010) was subsequently applied to each sample, and columns were incubated on ice for 15 minutes. After incubation, samples were washed three additional times before RNA was eluted in RNase-free water. RNA concentration was then measured, and the samples were stored at -80°C. subsequently used for qPCR and RNA sequencing.

### qPCR

Reverse transcription was carried out with iScript cDNA Synthesis Kit (BIO-RAD) in a 20 μl reaction system (4 μl 5x iScript Reaction Mix, 1 μl iScript Reverse Transcriptase, 1 μg RNA, Nuclease-free water to 20 μl) to obtain cDNA. The reaction conditions was as follows: Priming 5 min at 25°C; Reverse transcription 20 min at 46°C; RT inactivation 1 min at 95°C; Hold at 4°C (Optional step). Primers for real-time quantitative reverse transcription PCR (qRT-PCR) were designed and obtain from Millipore Sigma. qRT-PCR was performed using SYBR Green Universal Master Mix (Thermo Scientific) in QuantStudio 3 (Thermo Scientific Each 20 μL reaction contained 10 μL of SYBR Green Mix, 1 μL of cDNA template, 0.5 μL each of forward and reverse primers, and 8 μL of nuclease-free water. Amplification was conducted using a standard cycling program with a melting curve analysis. For each condition, three biological replicates and three technical replicates were analyzed. Data were processed using quickQrtPCR/TBtools-II and calculated using the 2^−ΔΔCt^ method.

### mRNA-sequencing and Bioinformatic Analysis

RNA sequencing libraries (unstranded) were prepared using the NEBNext Ultra II RNA Library Prep Kit for Illumina using manufacturer’s instructions (NEB, Ipswich, MA, USA). Briefly, mRNAs were initially enriched with Oligod(T) beads. Enriched mRNAs were fragmented for 15 minutes at 94 °C. First strand and second strand cDNA were subsequently synthesized. cDNA fragments were end repaired and adenylated at 3’ends, and universal adapters were ligated to cDNA fragments, followed by index addition and library enrichment by PCR with limited cycles. The sequencing library was validated on the Agilent TapeStation (Agilent Technologies, Palo Alto, CA, USA), and quantified by using Qubit 3.0 Fluorometer (Invitrogen, Carlsbad, CA) as well as by quantitative PCR (KAPABiosystems, Wilmington, MA, USA).

The sequencing libraries were multiplexed and clustered onto a flowcell on the Illumina NovaSeq instrument according to manufacturer’s instructions. The samples were sequenced using a 2×150bp Paired End (PE) configuration with a goal of 50M reads per sample. Image analysis and base calling were conducted by the NovaSeq Control Software (NCS). Raw sequence data (.bcl files) generated from Illumina NovaSeq was converted into fastq files and de-multiplexed using Illumina bcl2fastq 2.20 software. One mis-match was allowed for index sequence identification.

Raw data are available through NCBI GEO (GSE329018). Data analysis was completed in Galaxy [1, 2]. Quality control and adapter trimming were completed with *Trim Galore!* [3], which is a wrapper around *Cutadapt* [4] and *FastQC* [5] (Read 1 sequence: AGATCGGAAGAGCACACGTCTGAACTCCAGTCAC; Read 2 sequence:

AGATCGGAAGAGCGTCGTGTAGGGAAAGAGTGT). Trimmed reads were aligned to the hg38 genome using *HISAT2* [6], gene expression was quantified using *featureCounts* [7], and differential gene expression was determined using *DESeq2* [8]. PCA plots were generated with DESeq2. Normalized gene expression values (FPKM) were determined with *Cufflinks* [9]. Annotations of upregulated genes were completed using DAVID [10] where upregulated genes (determined via *DESeq2*) had a log_2_ fold change greater than or equal to 1 and an adjusted p value less than 0.01. Heat map values were calculated as the log_2_ of the biological replicate’s FPKM value divided by the mean of all sample FPKM values for a given gene.

### Magic red assay

U2OS WT, p62-KO, and p62-KO cells stably expressing WT p62 (rescue) were seeded into four-chamber live cell imaging dishes. Cells were incubated with Magic Red (1: 250 dilution, Antibodies Inc, SKU:942) for 30 mins before first imaging. Subsequently, cells were treated with 500 µM LLOME 10 mins, followed by a second imaging session. A third imaging were performed after washing cells with prewarm DMEM thoroughly and 30-min incubation. Magic Red was maintained at the same concentration throughout. Live-cell imaging was conducted on a Leica SP8 LIGHTNING confocal system equipped with an Okolab stage-top incubator, with Magic Red fluorescence excited at 550 nm and emission collected at 610 nm.

### Identification of CASM effectors

To identify ATG8 effectors during lysosome membrane permeabilization, we filtered our lysosome proteomics dataset^3^ to retain candidate proteins showing >2-fold increased lysosomal recruitment upon LLOME treatment compared with control. We then queried these candidates against the BioGRID interaction database^5^ to identify proteins with evidence of direct interaction with ATG8-family proteins.

### iPSC-derived Sensory Neuron Differentiation and Immunofluorescence

ND2.0 human induced pluripotent stem cells (iPSCs) were differentiated into neural crest cells (NCCs) using the STEMdiff™ Neural Crest Differentiation Kit (#08610, STEMCELL Technologies). NCCs were subsequently specified into sensory neurons using the STEMdiff™ Sensory Neuron Differentiation Kit (#100-0341, STEMCELL Technologies) and matured in STEMdiff™ Sensory Neuron Maturation Medium (#100-681, STEMCELL Technologies).

To examine p62 recruitment in neurons, matured sensory neurons were treated with vehicle or 500 µM LLOMe for 60 min. Cells were fixed with 4% paraformaldehyde (PFA) in PBS for 20 min, permeabilized with 0.1% Triton X-100 in PBS for 2 min, and blocked with 1x blocking buffer (Thermo Fisher Scientific, #37565) for 1 h at room temperature. Samples were then incubated with primary antibody against MAP2 (1:5000, #ab5392, Abcam), p62 (1:500, #sc-28359, Santa Cruz) and LAMP1 (1:500, #9091, Cell singaling technology) at 4°C overnight. After three PBS washes, cells were incubated with secondary antibodies for 1 h at room temperature. Following final washes, coverslips were mounted using VECTASHIELD HardSet Antifade Mounting Medium (Vector Laboratories, #H-1700). Confocal imaging was performed on a Leica SP8 LIGHTNING system using Leica Application Suite X (v3.5.5.19976) software.

### Mass spectrometry and data processing

Samples were separated ∼1.5 cm on a 10% Bis-Tris Novex mini-gel (Invitrogen) using the MES buffer system. The gel was stained with coomassie and excised into ten equally sized segments. Gel segments were processed using a robot (DigestPro, CEM) with the following protocol: Washed with 25mM ammonium bicarbonate followed by acetonitrile. Reduced with 10mM dithiothreitol at 60°C followed by alkylation with 50mM iodoacetamide at RT. Digested with trypsin (Promega) at 37°C for 4h. Quenched with formic acid and the supernatant was analyzed directly without further processing.

The gel digests were analyzed by nano LC/MS/MS with a Waters M-class HPLC system interfaced to a ThermoFisher Fusion Lumos. Peptides were loaded on a trapping column and eluted over a 75 μm analytical column at 350nL/min; both columns were packed with XSelect CSH C18 resin (Waters). A 30min gradient was employed. The mass spectrometer was operated in data-dependent mode, with MS and MS/MS performed in the Orbitrap at 60,000 FWHM resolution and 15,000 FWHM resolution, respectively. APD was turned on. The instrument was run with a 3s cycle for MS and MS/MS.

Data were searched using a local copy of Mascot with the following parameters: Enzyme: Trypsin; Database: Swissprot Human (concatenated forward and reverse plus common contaminants); Fixed modification: Carbamidomethyl (C); Variable modifications: Oxidation (M), Acetyl (Protein N-term), Deamidation (NQ), Pyro-Glu (N-term Q); Mass values: Monoisotopic; Peptide Mass Tolerance: 10 ppm; Fragment Mass Tolerance: 20 ppm; Max Missed Cleavages: 2. Mascot DAT files were parsed into the Scaffold software for validation, filtering and to create a nonredundant list per sample. Data were filtered at 1% protein and peptide level FDR and requiring at least two unique peptides per protein.

### Software

Confocal images were captured using a Leica SP8 LIGHTNING confocal system, utilizing the Leica Application Suite X 3.5.5.19976 software. The images were uniformly processed and compiled in Adobe Photoshop 21.0.2.

A Python program was used to quantify colocalization images captured with Confocal Microscopy. For the quantification of fluorescence images, the outlines of randomly selected cells were manually annotated in each image by an investigator blinded to the allocation. In order to quantify the fraction of protein A with protein B recruitment in puncta, a threshold was implemented to minimize diffuse protein B signals. The thresholds for both proteins were set based on the percentile of the target protein’s signal intensity within an individual cell, with a minor constant adjustment. The same threshold settings were used for all cells from the same experiments. Accurate threshold applications were ensured by manual checking.

Following data acquisition, data comparisons for all groups were visualized in GraphPad Prism, which was also used to calculate p values. The full code and environment configuration can be found on GitHub and Zenodo. See Key Resources Table.

### Statistics and reproducibility

All experiments, except for the Lyso-TurboID mass spectrometry which was conducted once, were independently reproduced at least three times. Rigorous standards ensured the results were robust and unbiased. No statistical techniques were applied to decide the sample size beforehand. Investigators were kept blind to the allocation during the imaging and analysis phases. Statistical significance was determined by unpaired, two-tailed t-test with the Analysis ToolPak in Microsoft Office Excel 2016 or one/two-way ANOVA with Tukey’s multiple comparison test in GraphPad Prism 8.0.2. Unless otherwise specified, data are presented as Mean ± SEM; N numbers are defined in the figure legends; when applicable, the total number of cells quantified was pooled from three independent experiments.

---

**Figure S1.**
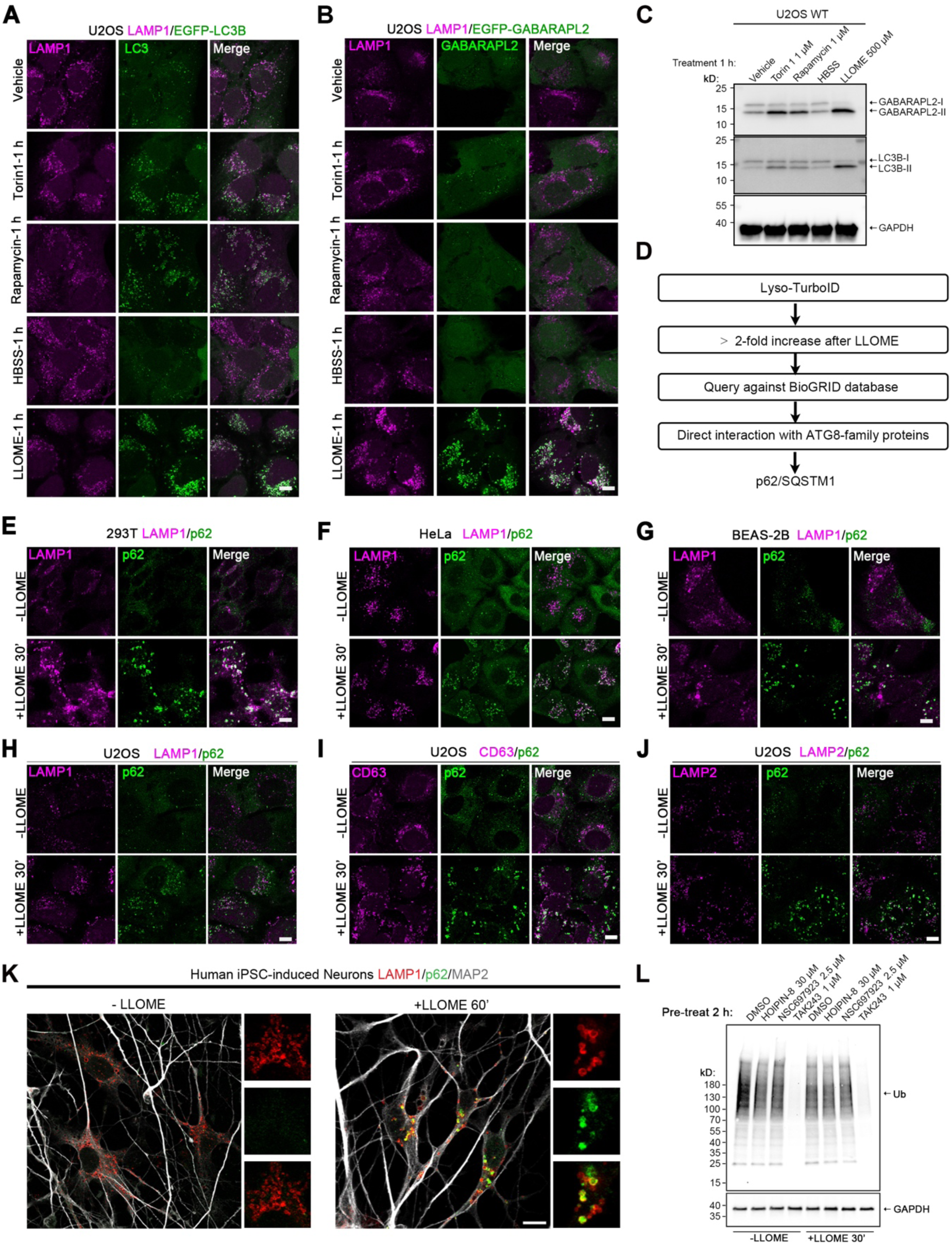
LMP triggers strong ATG8ylation and lysosomal recruitment of p62, related to Figure 1. (A) Lysosomal damage by LLOME induces stronger LC3B puncta compared with autophagy-inducing stressors. U2OS cells stably expressing EGFP-LC3B were treated as indicated for 1 h, followed by fixation and immunostaining of endogenous LAMP1. (B) Lysosomal damage by LLOME induces stronger GABARAPL2 puncta compared with autophagy-inducing stressors. U2OS cells stably expressing EGFP-GABARAPL2 were treated as indicated for 1 h, followed by fixation and immunostaining of endogenous LAMP1. (C) Immunoblot showing stronger ATG8ylation following LLOME treatment compared with autophagy-inducing stressors. (D) Bioinformatics analysis of Lyso-TurboID hits identifies p62/SQSTM1 as a top ATG8-interacting proteins enriched on damaged lysosomes. (E-H) Endogenous p62 recruitment to damaged lysosomes in different cell types including 293T (E), HeLa (F), and BEAS-2B (G), and U2OS (H). (I-J) Further examination of p62 recruitment by co-staining p62 with additional lysosomal markers, CD63 (I) and LAMP2 (J). (K) Representative images showing p62 recruitment to lysosomes in human iPSC-derived neurons treated with 500 µM LLOMe for 1h. (L) Immunoblot showing the blockage of ubiquitination by different inhibitors. HOIPIN-8 specifically inhibits HOIP (HOIL-1–interacting protein) to block linear (M1-linked) ubiquitin chain formation; NSC697923 targets the E2 enzyme UBE2N (Ubc13) to inhibiting K63-linked ubiquitination; TAK-243 inhibits the E1 activating enzyme UBA1, leading to a global shutdown of ubiquitin conjugation. Bar, 10 μm.

**Figure S2.**
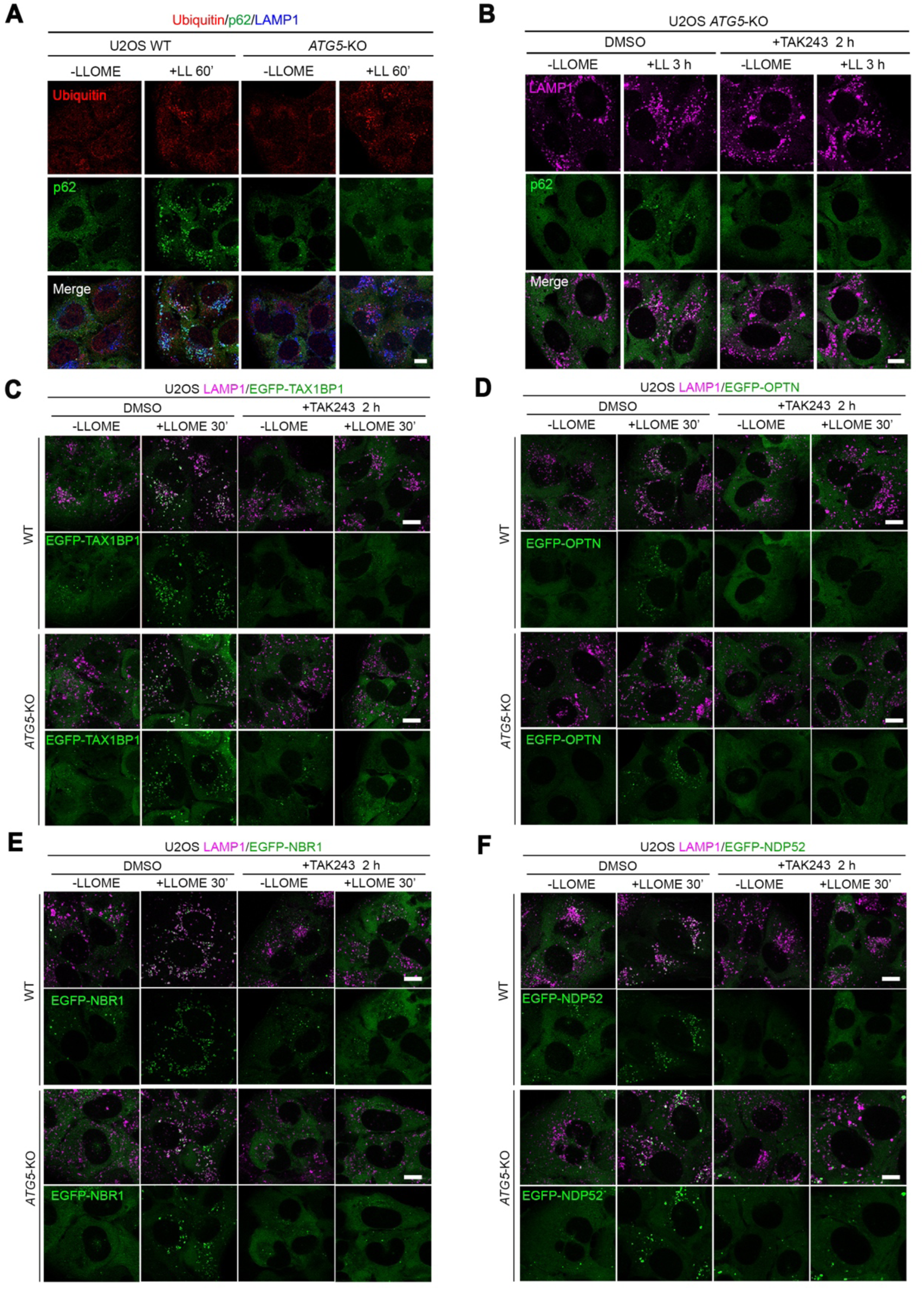
Ubiquitination plays a minor role for lysosomal p62 recruitment but is a major mechanism for recruiting other autophagy adaptors to damaged lysosomes, related to Figure 1. (A) *ATG5* knockout does not affect the levels of LMP-induced ubiquitin puncta. U2OS WT or *ATG5*-KO cells were treated with vehicle or 500 μM of LLOME for indicated time, followed by fixation and immunostaining of endogenous ubiquitin, p62 and LAMP1. (B) Ubiquitination mediates weak p62 recruitment after prolonged lysosomal damage in *ATG5*-KO cells. U2OS *ATG5*-KO cells were treated with DMSO or TAK243 for 2 h, and then treated with vehicle or 500 μM of LLOME for 3 h, followed by fixation and immunostaining of endogenous p62 and LAMP1. (C) The recruitment of autophagy adaptor TAX1BP1 to damaged lysosomes is dependent on ubiquitination but not ATG5. U2OS WT or *ATG5*-KO cells stably expressing EGFP-TAX1BP1 were treated with DMSO or TAK243 for 2 h, and then treated with vehicle or 500 μM of LLOME for 30 min, followed by fixation and immunostaining of endogenous LAMP1. (D) The recruitment of autophagy adaptor OPTN to damaged lysosomes is dependent on ubiquitination but not ATG5. U2OS WT or *ATG5*-KO cells stably expressing EGFP-OPTN were treated with DMSO or TAK243 for 2 h, and then treated with vehicle or 500 μM of LLOME for 30 min, followed by fixation and immunostaining of endogenous LAMP1. (E) The recruitment of autophagy adaptor NBR1 to damaged lysosomes is dependent on ubiquitination but not ATG5. U2OS WT or *ATG5*-KO cells stably expressing EGFP-NBR1 were treated with DMSO or TAK243 for 2 h, and then treated with vehicle or 500 μM of LLOME for 30 min, followed by fixation and immunostaining of endogenous LAMP1. (F) The recruitment of autophagy adaptor NDP52 to damaged lysosomes is dependent on ubiquitination but not ATG5. U2OS WT or *ATG5*-KO cells stably expressing EGFP-NDP52 were treated with DMSO or TAK243 for 2 h, and then treated with vehicle or 500 μM of LLOME for 30 min, followed by fixation and immunostaining of endogenous LAMP1. Bar, 10 μm.

**Figure S3.**
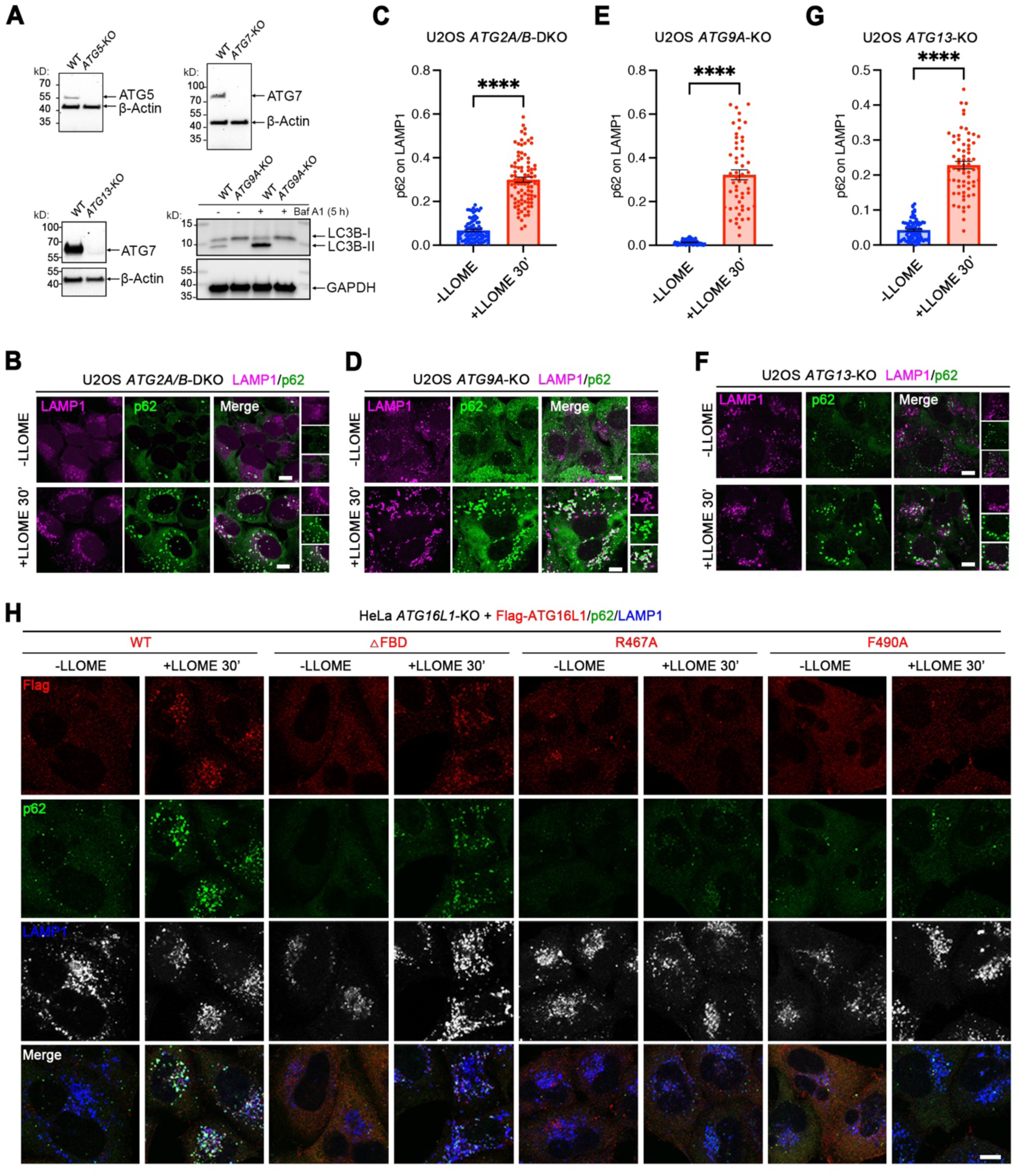
p62 recruitment to damaged lysosomes is dependent on CASM but not macroautophagy, related to Figure 2. (A) Immunoblot showing the knockout of *ATG5/7/13/9*. Note that we were not able to detect the ATG9A band, but the *ATG9A*-KO cells showed a complete loss of LC3B lipidation. (B) *ATG2A/B* double knockout (DKO) does not block p62 recruitment to damaged lysosomes. U2OS *ATG2A/B*-DKO cells^14^ were treated with vehicle or 500 μM of LLOME for 30 min, followed by fixation and immunostaining of endogenous p62 and LAMP1. (C) Quantification of p62 recruitment to lysosomes in (B). Data are means ± SEM; n = 80 and 91 random cells, respectively. (D) *ATG9A* knockout does not block p62 recruitment to damaged lysosomes. U2OS *ATG9A*-KO cells were treated with vehicle or 500 μM of LLOME for 30 min, followed by fixation and immunostaining of endogenous p62 and LAMP1. (E) Quantification of p62 recruitment to lysosomes in (D). Data are means ± SEM; n = 68 and 53 random cells, respectively. (F) *ATG13* knockout does not block p62 recruitment to damaged lysosomes. U2OS *ATG13*-KO cells were treated with vehicle or 500 μM of LLOME for 30 min, followed by fixation and immunostaining of endogenous p62 and LAMP1. (G) Quantification of p62 recruitment to lysosomes in (F). Data are means ± SEM; n = 83 and 67 random cells, respectively. (H) ATG16L1 recruitment to damaged lysosomes during LMP is dependent on its WD40 domain rather than the FBD domain. HeLa *ATG16L1*-KO cells re-expressing Flag-ATG16L1 or its mutants were treated with vehicle or 500 μM of LLOME for 30 min, followed by fixation and immunostaining of Flag-tag and endogenous p62 and LAMP1. Data were analyzed by Student’s t test for (C), (E) and (G). WT, wild type. KO, knockout. Bar, 10 μm.

**Figure S4.**
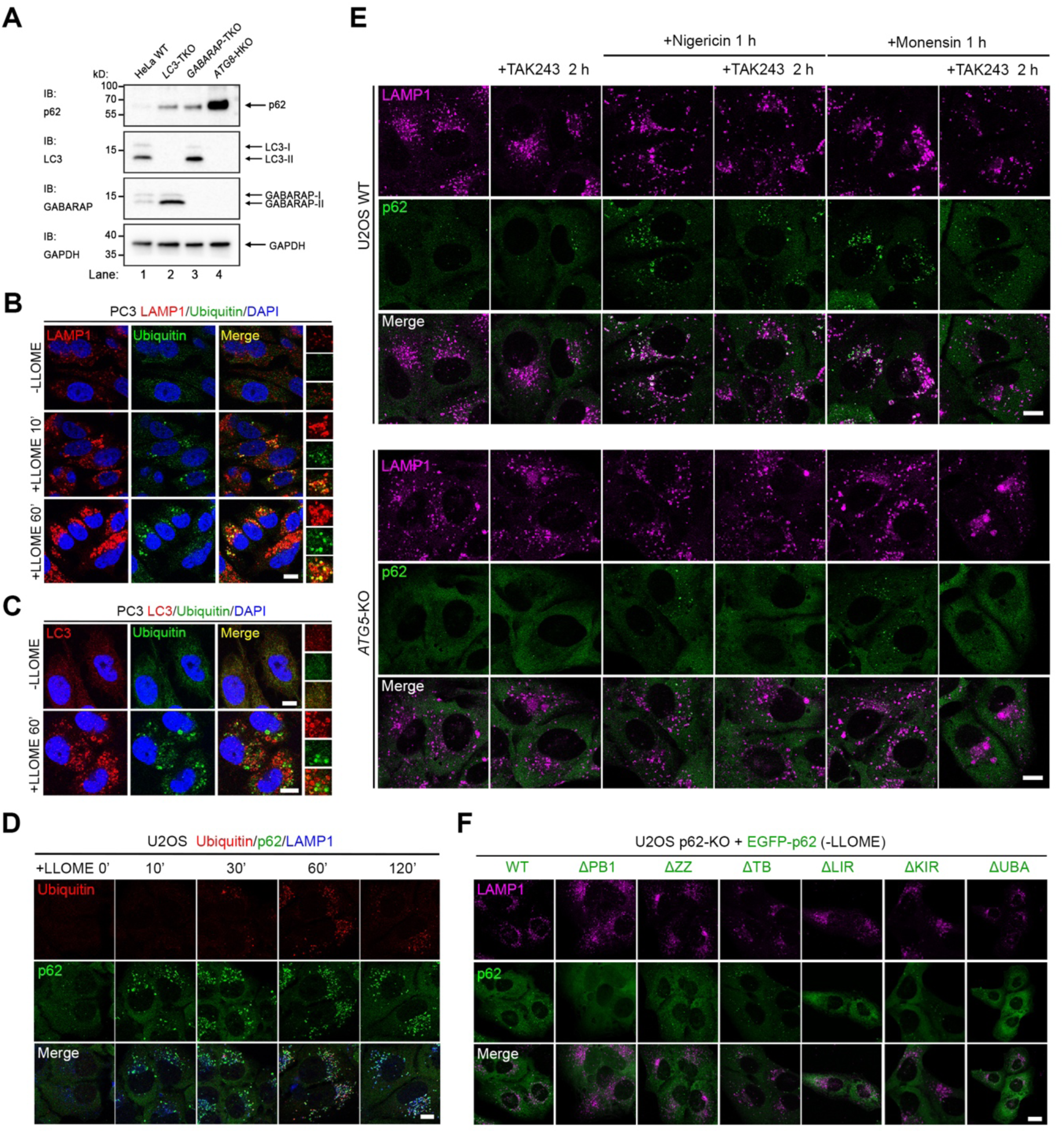
ATG8ylation rather than ubiquitination recruits p62 to damaged lysosomes, related to Figure 3. (A) Immunoblot showing the knockout of ATG8 family proteins in HeLa cells. (B) Endogenous ubiquitin recruitment to a small fraction of damaged lysosomes. PC3 cells were treated with vehicle or 500 μM of LLOME as indicated, followed by fixation and immunostaining of endogenous ubiquitin and LAMP1. (C) LC3 and ubiquitin recruitment to different compartments. PC3 cells were treated with vehicle or 500 μM of LLOME for 1 h, followed by fixation and immunostaining of endogenous ubiquitin and LC3. (D) Earlier lysosomal recruitment of p62 than ubiquitin following LLOME treatment. U2OS cells were treated with vehicle or 500 μM of LLOME as indicated, followed by fixation and immunostaining of endogenous ubiquitin, p62 and LAMP1. (E) Nigericin and monensin-induced p62 recruitment to damaged lysosomes largely depend on ATG5 rather than ubiquitination. U2OS cells were pre-treated with DMSO or TAK243 for 2 h and then treated with vehicle or 100 nM nigericin or 18.5 μM monensin for 1 h, followed by fixation and immunostaining of endogenous p62 and LAMP1. (F) The localizations of EGFP-p62 and mutants stably expressed in U2OS p62-KO cells in resting conditions. Representative images from vehicle treated cells for the experiment in **Figure 3G**. Quantified in **Figure 3H**. KO, knockout. Bar, 10 μm.

**Figure S5.**
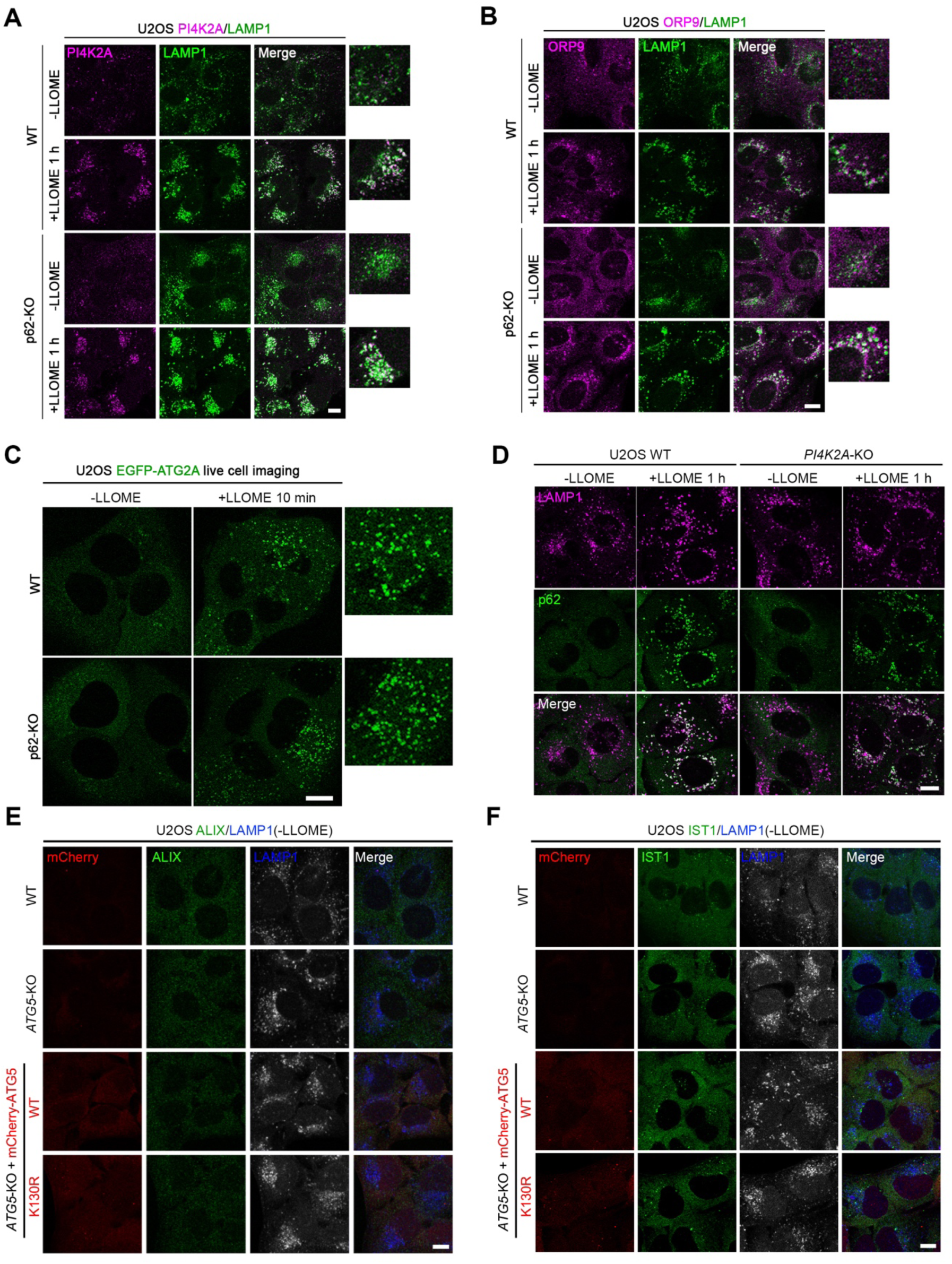
p62 recruitment is independent from PITT pathway, related to Figure 4. (A) p62-KO does not affect PI4K2A recruitment to damaged lysosomes. U2OS cells were treated with vehicle or 500 μM of LLOME for 1 h, followed by fixation and immunostaining of endogenous PI4K2A and LAMP1. (B) p62-KO does not affect ORP9 recruitment to damaged lysosomes. U2OS cells were treated with vehicle or 500 μM of LLOME for 1 h, followed by fixation and immunostaining of endogenous ORP9 and LAMP1. (C) p62-KO does not affect EGFP-ATG2A puncta formation in response to lysosomal damage. U2OS cells stably expressing EGFP-ATG2A were treated with vehicle or 500 μM of LLOME, followed by live cell imaging. (D) *PI4K2A*-KO does not affect p62 recruitment to damaged lysosomes. U2OS cells were treated with vehicle or 500 μM of LLOME for 1 h, followed by fixation and immunostaining of endogenous p62 and LAMP1. (E,F) The localizations of ALIX (E) and IST1 (F) in resting conditions under different genetic background. Bar, 10 μm.

**Figure S6.**
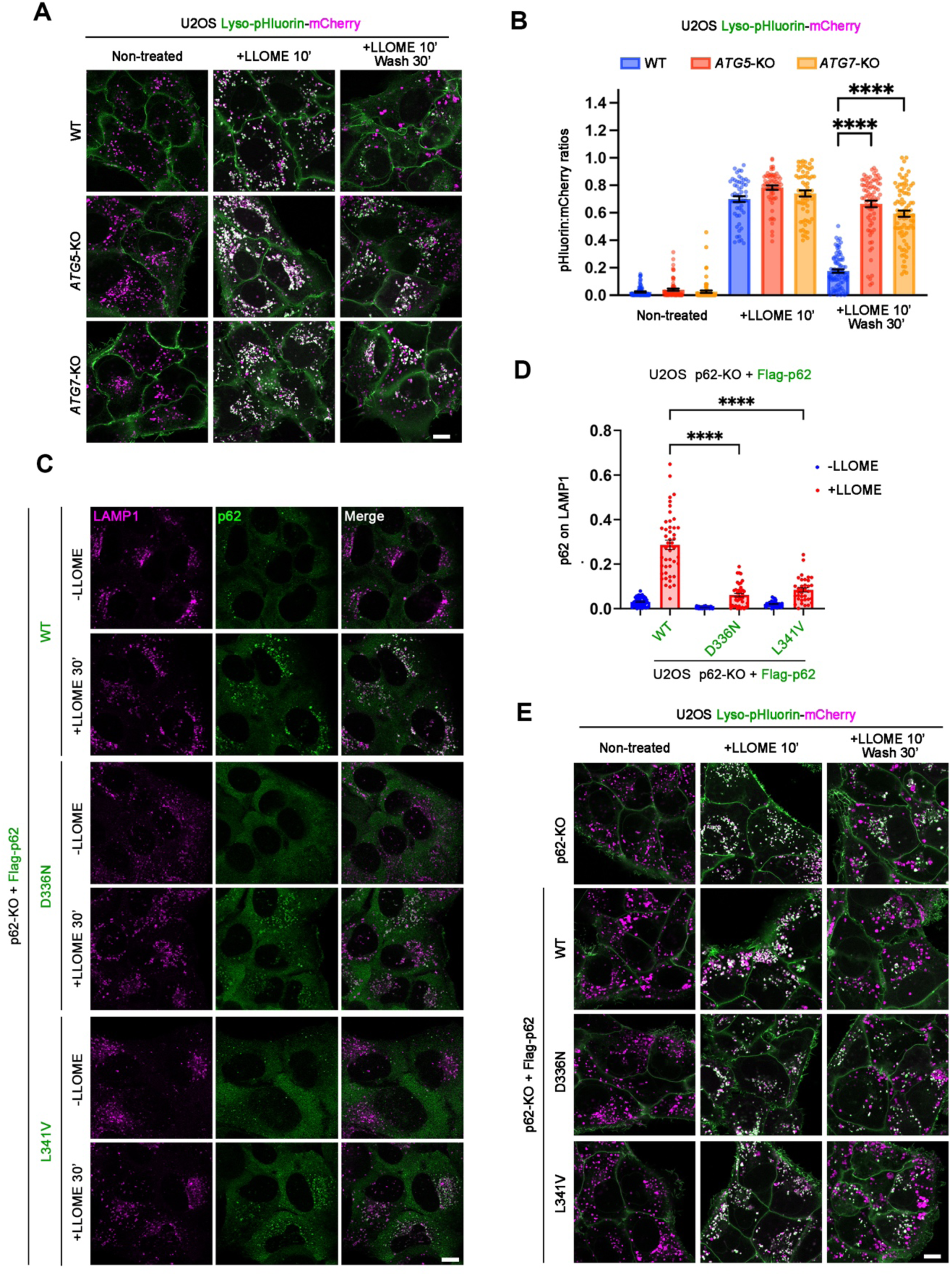
The CASM-p62 axis recruits KEAP1 to damaged lysosomes, related to Figure 5. (A) Lyso-pHluorin-mCherry images showing lysosomal repair defects in *ATG5*-KO and *ATG7*-KO cells. WT and the KO cells stably expressing Lyso-pHluorin-mCherry were treated with 500 μM LLOME as indicated, followed by live cell imaging. (B) Quantification of the pHluorin-to-mCherry signal ratios in (A). Data are means ± SEM; n = 84, 69, 75, 55, 62, 62, 87, 79, 84 random cells, respectively. (C) Disease mutants of p62, D336N and L341N, impair its recruitment to damage lysosomes. U2OS p62-KO cells stably expressing Flag-tagged p62 WT or mutants were treated with 500 μM LLOME as indicated, followed by fixation and immunostaining of endogenous p62 and LAMP1. (D) Quantification of p62 signals on LAMP1 in (C). Data are means ± SEM; n = 51, 33, 32, 45, 39, 35 random cells, respectively. (E) Lyso-pHluorin-mCherry images showing lysosomal repair defects of p62 D334N and L341N mutants. Cells stably expressing Lyso-pHluorin-mCherry were treated with 500 μM LLOME as indicated, followed by live cell imaging. Data were analyzed by two-way ANOVA with Tukey’s multiple comparison tests for (B) and (D). WT, wild type. KO, knockout. Bar, 10 μm.

**Figure S7.**
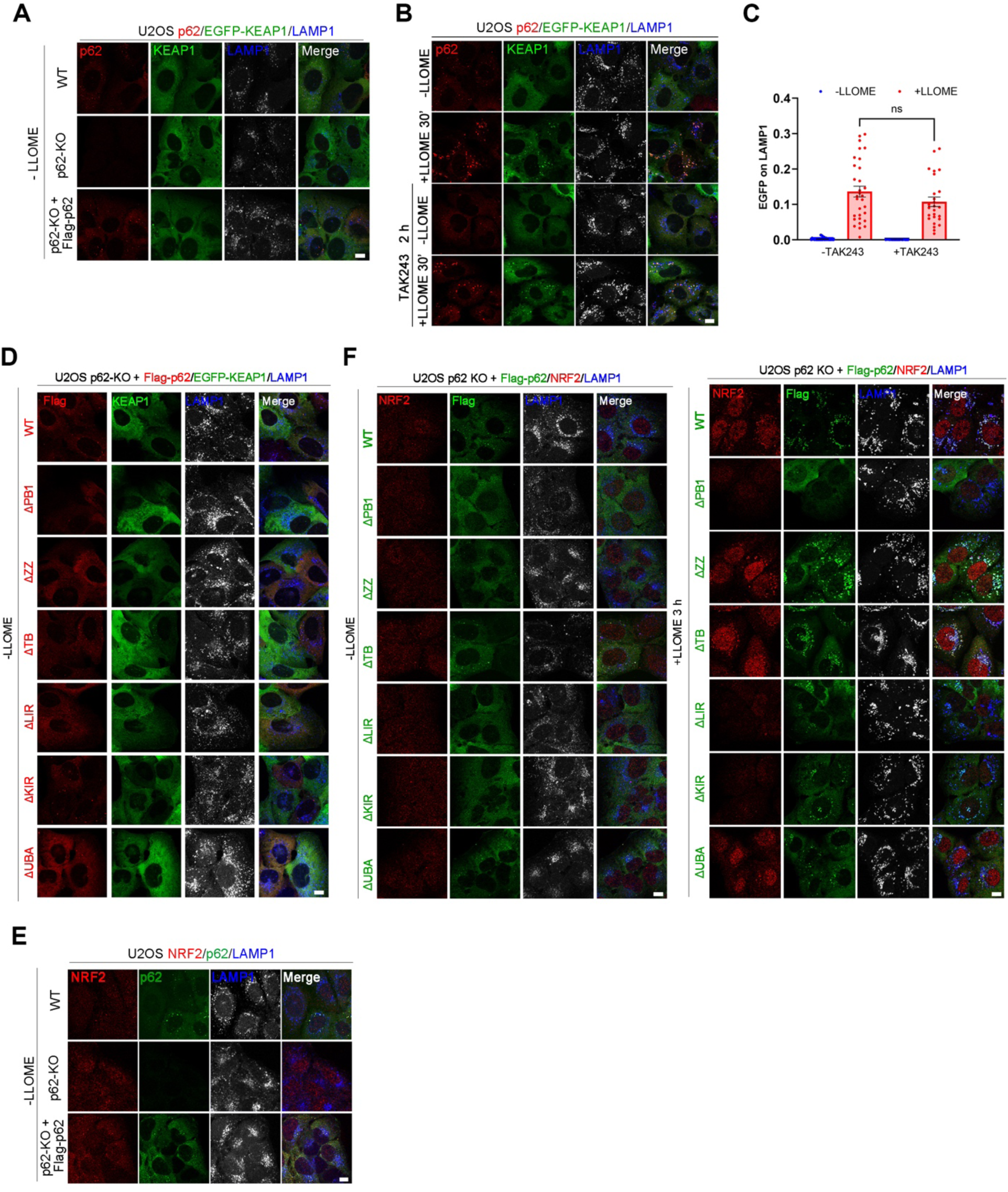
Nrf2 mediates CASM-p62-KEAP1 axis-triggered redox defense, related to Figure 6 and 7. (A) EGFP-KEAP1 is evenly distributed in the cytosol under resting conditions in the indicated cell lines. Negative control for Figure 6C. Quantified in Figure 6D. (B) Inhibiting total cellular ubiquitination by TAK243 (1 μM 2 h pretreatment) does not affect LLOME-induced KEAP1 recruitment to lysosome. U2OS cells stably expressing EGFP-KEAP1 were treated as indicated, followed by fixation and immunostaining of endogenous p62 and LAMP1. (C) Quantification of EGFP-KEAP1 recruitment to lysosomes in (B). Data are means ± SEM; n = 33, 31, 23, 26 random cells, respectively. Data were analyzed by two-way analysis of variance (ANOVA) with Tukey’s multiple comparison tests. (D) EGFP-KEAP1 is evenly distributed in the cytosol under resting conditions in the indicated cell lines. Negative control for Fig 6G. Quantified in Fig 6H. (E) NRF2 subcellular localization under resting conditions in the indicated cell lines. Negative control for Fig 7A. Quantified in Fig 7B. (F) Lysosomal recruitment of p62 promotes LMP-induced NRF2 nuclear translocation. Quantified in Fig 7E. U2OS p62-KO cells stably expressing Flag-p62 WT or indicated mutants were treated with or without 500 μM LLOME for 3 h, followed by fixation and immunostaining of Flag-p62 and endogenous NRF2. WT, wild type. KO, knockout. Bar, 10 μm.

**Figure S8.**
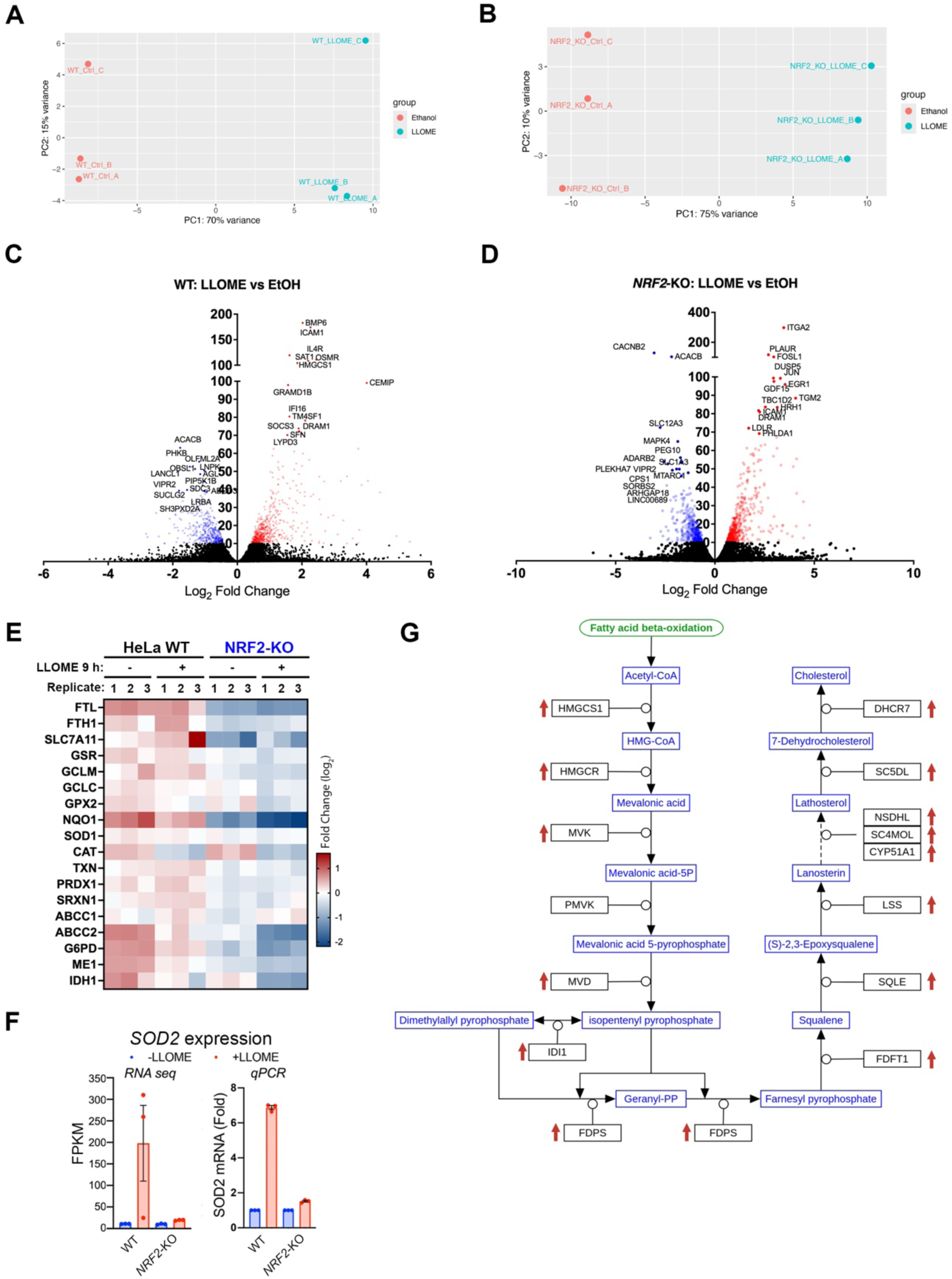
NRF2 mediates CASM-p62-KEAP1 axis-triggered redox defense, related to Figure 7. (A) Principal Component Analysis (PCA) plot of RNA-seq data in HeLa wild type cells. Each point represents a sample, and the colors correspond to different experimental groups. (B) PCA plot of RNA-seq data in HeLa *NRF2*-KO cells. Each point represents a sample, and the colors correspond to different experimental groups. (C) Volcano plot depicting differential expression of genes in WT HeLa cells after 9 h of LLOME (500 μM) treatment. (D) Volcano plot depicting differential expression of genes in *NRF2*-KO HeLa cells after 9 h of LLOME (500 μM) treatment. (E) Heatmap displaying the expression pattern of canonical NRF2 target genes in WT and *NRF2*-KO HeLa cells. (F) RNA seq and q-PCR results showing that lysosomal damage by LLOME triggers NRF2-dependent upregulation of *SOD2*, which encodes a mitochondrial antioxidative enzyme. (G) Illustration of genes from the cholesterol biosynthesis pathway (https://pathway-viewer.toolforge.org/embed/WP197) that are upregulated in a NRF2-dependent manner in response to lysosomal damage by LLOME.

